# A systems-level view of nucleus pulposus mechanotransduction and its regulatory control

**DOI:** 10.64898/2026.11.20.768129

**Authors:** Zerihun G. Workineh, Francis Kiptengwer Chemorion, Jérôme Noailly

**Affiliations:** Barcelona Centre for New Medical Technologies (BCN MedTech), Department of Engineering, Universitat Pompeu Fabra, Barcelona, 08018, Spain

## Abstract

Intervertebral disc degeneration (IDD), a major contributor to chronic low back pain, is strongly influenced by mechanical loading- dependent regulation of nucleus pulposus (NP) cell phenotype. Previous studies have characterized individual mechanosensors and specific signalling pathways involved in NP mechanotransduction. However, the regulatory interactions through which these components integrate mechanical cues into coordinated cellular responses remain incompletely understood. Here, we present an NP-specific, signed, directed regulatory network model (RNM) comprising 146 molecular nodes and 353 literature-supported interactions. The network connects mechanosensors with signalling cascades, metabolic and redox regulators, transcription factors, extracellular matrix (ECM) effectors, inflammatory mediators, and cell-fate pathways. Using the SQUADS semi-quantitative formalism, which accommodates limited kinetic parameter availability, we simulated steady-state responses to hypo-loading (Hypo), normal loading (NL), and hyper-loading (HL). We also analyzed network topology and screened 48 single- and dual-node perturbations under sustained HL. NL maintained high activity of the ECM-associated regulators ACAN, COL2A1, SOX9, and TIMP3, supported by AKT1, FAK, and antioxidant regulation. HL promoted inflammatory and catabolic regulation through mediators such as NF-*κ*B, AP-1, MMP13, and ADAMTS4/5, accompanied by increased ROS, BAX, and CASP3 activity. Hypo reduced anabolic activity while retaining calcium, ROS, AMPK, and apoptotic signalling, suggesting a possible adhesion-dependent stress mechanism. Model falsification against 39 literature-derived molecular directionality benchmarks yielded 37 concordant predictions (94.9%), providing indirect, direction-level support for the RNM. HSF1 and CCN2 showed contradictory responses. Simulated regulatory recovery was module-specific. SOX9 activation achieved 82.8% ECM-profile recovery, increasing to 85.9% with ROS suppression. SOX9 activation combined with NF-*κ*B suppression achieved 57.0% inflammatory/proteolytic-module recovery. FOXO activation with ROS suppression achieved 68.9% oxidative-stress/proteostasis-module recovery. NRF2 activation with PIEZO1 suppression achieved 37.5% cell-survival-module recovery. These predictions identify candidate molecular targets and intervention combinations for anabolic reinforcement, inflammatory suppression, redox protection, and mechanosensory modulation. The RNM provides a mechanistic framework for computational prioritization of candidate therapeutic targets and intervention combinations. Its predictions can guide subsequent preclinical validation in NP-cell and disc-explant models under controlled mechanical loading. Experimental validation, incorporation of time-dependent loading, and calibration of regulatory parameters are necessary to establish the therapeutic relevance of these predictions and improve the model’s translational applicability.

## 1 Introduction

Intervertebral disc (IVD) degeneration (IDD) is a major contributor to chronic low back pain, affecting more than half a billion people worldwide and representing a leading cause of disability and socioeconomic burden^1,2^. IDD is progressive and multifactorial, characterized by extracellular matrix (ECM) degradation, chronic inflammation, altered hydration, aberrant vascular and neural ingrowth, and loss of mechanical function^3–8^. These pathological features arise from the disruption of the specialized structure–function interactions within the IVD, involving the nucleus pulposus (NP), the annulus fibrosus (AF), and the cartilage endplate (CEP)^9–12^. Among these disc tissues, the NP plays a central role, as it maintains disc hydration and compressive load-bearing capacity through its proteoglycan-rich matrix, the breakdown of which being an early and critical degenerative event^6,13,14^.

Mechanical loading (ML) is an important determinant of intervertebral disc health and disease throughout life^15–20^. Under normal conditions, ML helps to maintain matrix organization, fluid pressurization, load bearing capacity, and disc height, and controlled deformations are thought to support functional solute transport through the cartilage endplate between the disc, that is avascular, and the vascularized vertebra^21,22^. In contrast, altered ML patterns that arise from life habits, injury, or previous impairment of disc biology and tissue turnover, can accelerate IDD^18,23–25^. At the cellular level, mechanical cues regulate NP cell metabolism and phenotype through mechanotransduction signaling pathways that control ECM synthesis, pro-inflammatory activity, oxidative stress, and cell survival^18,26–30^. Indeed, NP cells need ML, and reduced mechanical stimulation suppresses anabolic programs and impairs matrix maintenance^31^. In turn, excessive ML induces cytoskeletal strain, calcium dysregulation, oxidative stress, and upregulation of inflammatory mediators and matrix-degrading enzymes^32–36^.

Mechanotransduction refers to the process by which cells sense and convert mechanical stimuli into biochemical signals that, in turn, regulate cellular behavior per-se. Mechanical cues are primarily sensed through adhesion receptors, such as integrins (e.g., *α*5*β* 1, *α*V*β* 3), and through mechanosensitive ion channels (e.g., PIEZO1 and TRPV4)^35,37–41^. These inputs are transmitted through interconnected intra-cellular signalling cascades, including RhoA–ROCK–YAP/TAZ, MAPK–AP-1, PI3K–AKT–mTOR–HIF-1*α*, and Ca^2+^-dependent, pathways that integrate through metabolic, redox, and osmoadaptive regulators such as NRF2 and TonEBP/NFAT5^42–46^. Increasing evidence further indicates that mechanosensitive pathways can amplify pro-inflammatory cell activity and stress-associated programs under pathological ML^39,47–50^. Despite increasingly detailed molecular characterizations, extensive crosstalks and nonlinear feedback in these systems of signalling cascades make it difficult to determine how mechanical signals effectively integrate into coherent and stable phenotypic states^51–54^.

The unique microenvironment of the IVD makes it particularly challenging to understand the functional turnover of the organ, starting with the avascular nature and low-cellularity of the disc that tightly couple cell phenotypes with nutrient availability and waste removal^21,55–58^. Because disc cell nutrition depends predominantly on the transport of mass through the cartilage endplate and peripheral annulus, mechanical loading simultaneously alters fluid flow, solute gradients, and cellular energy demand, complicating the isolation of direct mechanotransductive effects from transport- and metabolism-driven regulation^21,59^. Furthermore, NP and AF responses are highly sensitive to the modality, amplitude, frequency, and duration, of mechanical loads, and experimental systems employ different strategies as per the ML per-se, and as per the preservation of native cell–matrix interactions, which contributes to heterogeneous observations^18,60–63^. Consequently, despite extensive characterization of mechanosensors and downstream pathways, it remains difficult to determine how their context-dependent crosstalk collectively gives rise to stable NP-cell states under distinct loading regimes.

Addressing this challenge motivates a systems-level framework able to integrate fragmented mechanobiological evidence and resolve emergent regulatory behavior emerging from multiple signal integration and feedback. Computational modeling of signaling networks has shown that cellular responses arise from the coordinated interaction of multiple pathways, making it difficult to infer system-level behavior from isolated signaling components alone^64–66^. Logical and regulatory network approaches enable systematic assembly of multiple molecular interactions and exploration of systems-level behaviors, in contexts where quantitative kinetic parameters remain incompletely defined^67,68^. For example, network-based models have predicted pathological phenotypes, allowing to identify key regulatory leverage points in cartilage homeostasis, degeneration, and inflammatory signaling^69–72^. Interactomes for the biochemical regulation of IVD NP cells were also proposed and successfully assessed, to integrate multiple downstream regulatory pathways, from cell membrane receptors activation, to effective pro-anabolic or pro-catabolic cell activity^72^. However, while mechanosensitive interactomes were proposed for articular cartilage chondrocyte^70^, an integrative regulatory framework that explicitly captures load-dependent mechanotransductive regulation in NP cells remains lacking.

In this study, we present a novel NP-specific, systems-level regulatory network model (RNM) of mechanotransduction. The curated network integrates more than 350 experimentally supported interactions across mechanosensors, intracellular signaling cascades, redox and inflammatory regulators, transcription factors, growth-factor pathways, and ECM effectors relevant to NP biology. Distinct mechanical loading regimes, hypo-, normal-, and hyper-loading, are represented as external regulatory inputs, enabling systematic interrogation of load-dependent network organization. We hypothesize that (H1) normal loading stabilizes a coordinated homeostatic phenotype through balanced activation of anabolic and protective programs, whereas (H2) deviations from this regime drive transitions toward alternative regulatory configurations characterized by loss of matrix maintenance, metabolic imbalance, and activation of stress-associated signaling. These hypotheses are examined through integrated analyses of network topology, dynamical behavior, falsification benchmarking, and perturbation screening to identify systems-level principles governing NP cell fate under mechanical loading. A complete list of gene symbols and their corresponding full names used throughout this manuscript is provided in Supplementary Material Table. S1.

## 2 Results

### 2.1 Network architecture and structural features

The curated NP-cell mechanotransduction network comprises 146 molecular nodes connected by 353 directed, signed regulatory interactions (Fig. 1). Nodes are colour-coded by functional class; mechanoreceptors, ion channels and related, kinase signalling modules, metabolic regulators, transcription factors, extracellular-matrix components, proteases, and stress-response mediators, providing a compact summary of the molecular scope captured by the model. The network exhibited an average node degree of 4.803, a diameter of 8.00, and a characteristic path length of 3.363. Network density was 0.033, with a clustering coefficient of 0.103. Network heterogeneity and centralization were 0.906 and 0.154, respectively.

**Figure 1.**
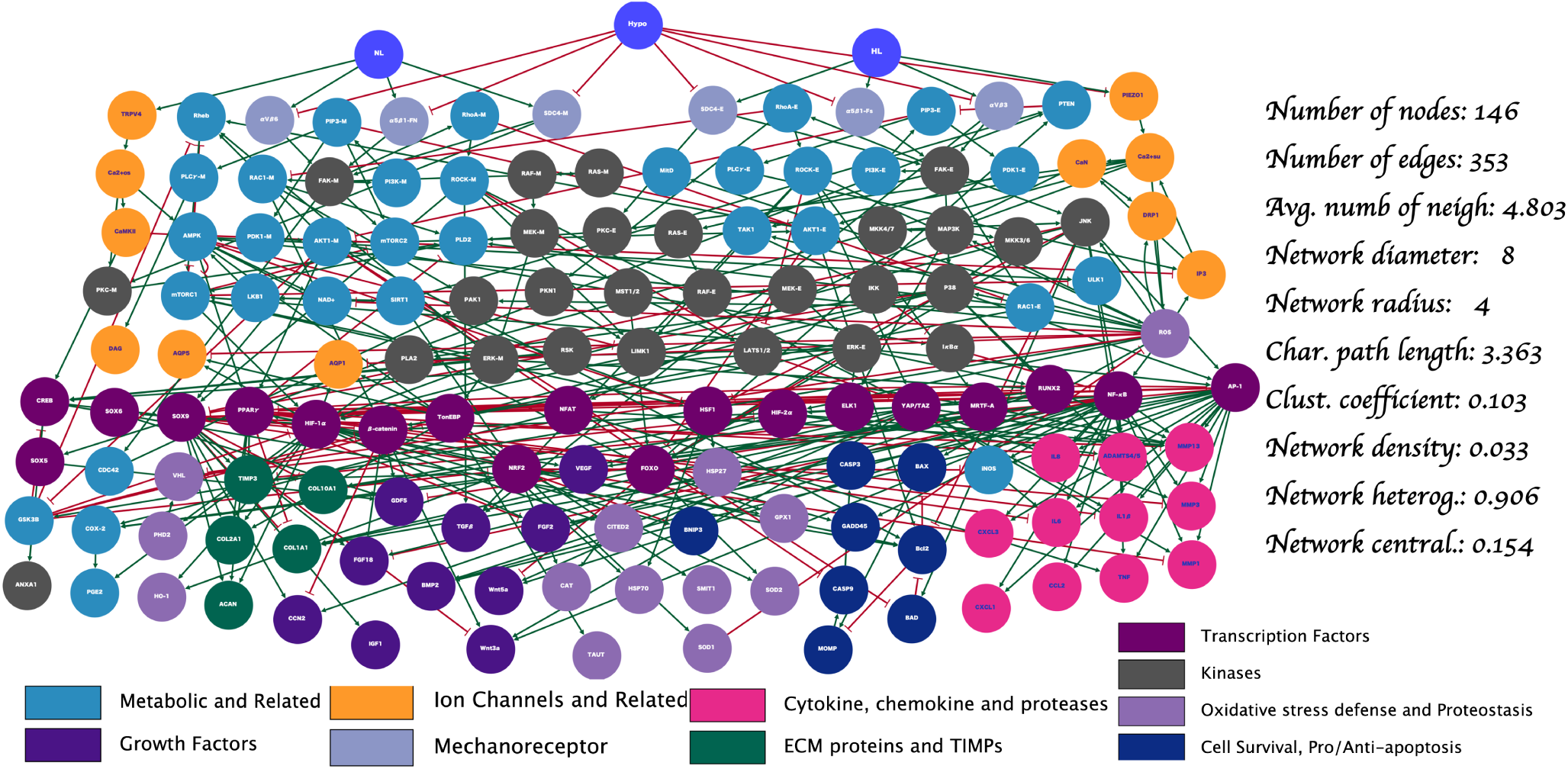
Curated mechanotransduction signalling network in NP cells, visualised using Cytoscape^73^. The network integrates 146 nodes and 353 signed regulatory interactions curated from peer-reviewed experimental studies. Nodes are colour-coded by functional category, and edges denote activating (green arrows) or inhibitory (red T-bars) regulation.

Signed out-degree analysis identified a subset of nodes with high outgoing regulatory influence (Fig. 2A). NF-*κ*B displayed the largest total out-degree (24 outgoing edges; 20 activating and 4 inhibitory), followed by SOX9, AP-1, ROS, and PPAR*γ*.

**Figure 2.**
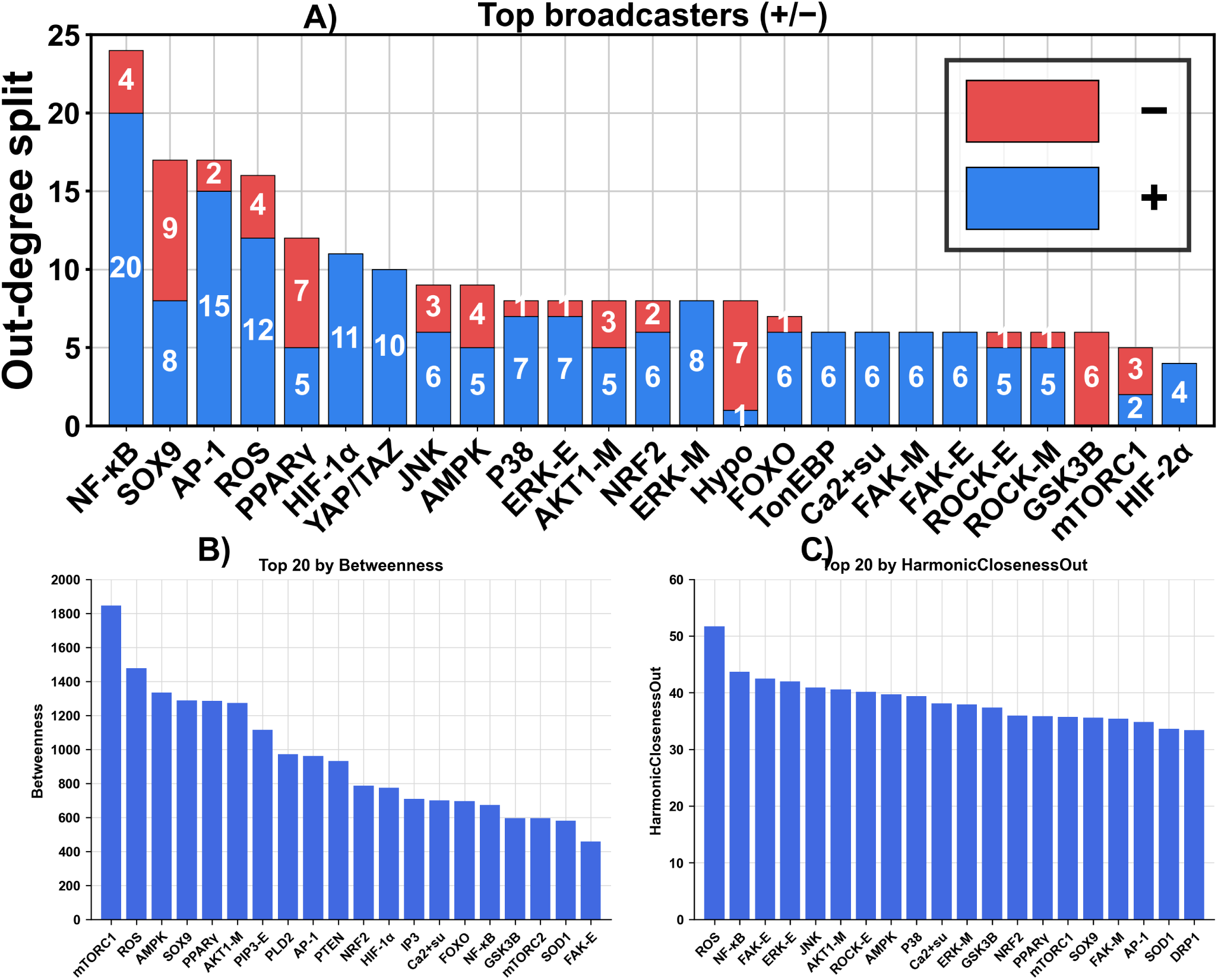
Topological metrics of the NP mechanotransduction network. (A) nodes ranked by signed out-degree, including the number of inhibitory (red) and activating (blue) outgoing edges, (B) nodes ranked by betweenness centrality, and (C) nodes ranked by harmonic closeness centrality.

Betweenness centrality highlighted nodes positioned as major topological bridges between network regions (Fig. 2B), with mTORC1 ranking highest, followed by ROS, AMPK, SOX9, PPAR*γ*, and additional signalling intermediates. Harmonic closeness centrality provided a complementary measure of global reachability in the directed network (Fig. 2C), ranking ROS highest, followed by NF-*κ*B, FAK-E, ERK-E and JNK.

### 2.2 Baseline steady-state activation profiles across mechanical loading regimes

Differences in final node activation among the three loading conditions were assessed using Kruskal–Wallis tests, followed by pairwise Wilcoxon rank-sum tests. The resulting *p*-values were adjusted for multiple comparisons using the Benjamini– Hochberg false discovery rate procedure^74^. An adjusted *q*-value below 0.05 was considered statistically significant. Matrix- associated readouts differed across all three loading regimes (Fig. 3A). *COL2A1*increased from 0.200 ± 0.340 under Hypo to 0.998 under NL, then decreased to 0.272 ± 0.438 under HL. *ACAN* and *TIMP3* likewise approached maximal activation under NL (0.999 and 0.998 ± 0.001, respectively), compared with 0.233 ± 0.376 and 0.000 under HL. The Hypo values of both markers were below their NL levels. In contrast, *COL1A1* and *COL10A1* were minimally active under NL (0.001 ± 0.001 and 0.000) and strongly active under HL (0.999 and 0.979 ± 0.033). Their Hypo activations were intermediate, with *COL10A1* at 0.449 ± 0.499. All corresponding NL–HL comparisons had adjusted *q <* 0.001. Growth-factor outputs showed additional differences (Fig. 3B). *IGF1* (1.000), *GDF5* (0.988), and *FGF18* (0.866) were strongly activated under NL but remained close to zero under Hypo. *CCN2* was also higher under NL (0.984) than HL. Conversely, HL favored *BMP2* (1.000), *FGF2* (0.990 ± 0.006), *Wnt3a* (0.990 ± 0.006), and *Wnt5a* (0.994 ± 0.008). Under Hypo, *VEGF*, *BMP2*, *FGF2*, and *Wnt3a* displayed intermediate activation, while *Wnt5a* remained low. *TGFβ* was active under both NL (0.846 ± 0.000) and HL (0.986 ± 0.000). The transcription-factor and inflammatory panels showed distinct profiles (Fig. 3C–D). Under NL, *SOX9* (0.999), *CREB* (0.994), *PPARγ* (0.902 ± 0.095), and *HIF-1α* (0.880 ± 0.095) showed high activation. Under HL, *NF-κB* (0.979 ± 0.033), *AP-1* (1.000), *RUNX2* (0.986), and *MRTF-A* (0.981 ± 0.031) were more active. Several HL-associated transcription factors, including *NF-κB*, *AP-1*, and *RUNX2*, had intermediate means under Hypo (0.41–0.49). *YAP/TAZ* remained active under NL (0.824) and HL (0.930), with little activity under Hypo. For inflammatory and matrix-degrading readouts, HL means approached one for *TNF*, *IL6*, *IL1β* , *MMP13*, and *ADAMTS4/5*, while their NL means were at or below 0.003 ± 0.003. Hypo means fell between these two conditions: approximately 0.440 ± 0.500 for *IL1β* , *IL6*, *TNF* and *ADAMTS4/5* (NL–HL, all *q <* 0.001).

**Figure 3.**
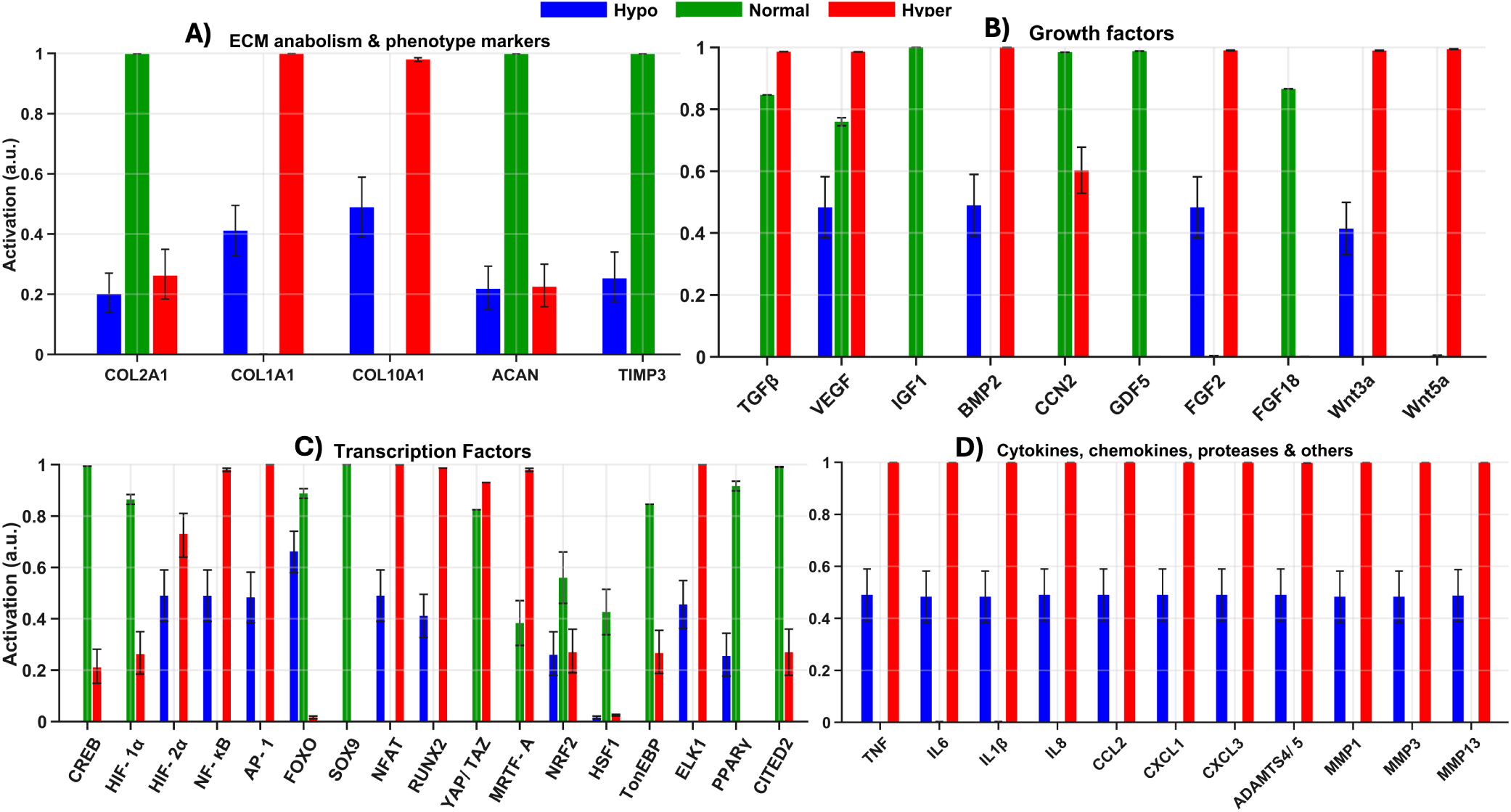
Baseline mean activation under Hypo (blue), NL (green), and HL (red). Bars represent means across 100 replicate simulations per condition; error bars indicate ±1 SD. (A) Extracellular-matrix and phenotype markers; (B) growth factors; (C) transcription factors; and (D) cytokines, chemokines, proteases, and related mediators. Activations are bounded between 0 and 1; symmetric SD whiskers can extend beyond these bounds.

Metabolic and calcium-associated nodes also differentiated Hypo from NL and HL (Fig. 4A–B). The *PI3K-M* (), *PIP3-M*, and *PDK1-M* nodes were nearly maximally active (1.000) under NL, as was *AKT1-M* (0.964 ± 0.035); these nodes were close to zero under HL and Hypo. Conversely, HL showed increased activation of *PI3K-E* (1.000), *PIP3-E* (0.722 ± 0.448), *PDK1-E* (0.720 ± 0.451), and *AKT1-E* (0.669 ± 0.419), compared with negligible activation under NL and Hypo. In contrast to those branches, *AMPK* and *ULK1* were appreciably active under Hypo (0.594 ± 0.381 and 0.703 ± 0.451, respectively). *PGE2* showed activity under all three loading regimes, including NL (0.833) and HL (1.000). The NL condition favored *Ca*^2+^, *CaMKII*, *PKC-M*, and *PLCγ-M* (each approximately 1.000), while HL favored 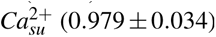 (0.979 ± 0.034), *PKC-E* (1.000), and *PLCγ-E* (0.720 ± 0.451). Under Hypo, the NL-associated calcium branch remained low, whereas 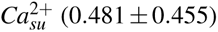 (0.481 ± 0.455) and *PKC-E* (0.477 ± 0.451) showed intermediate responses. *AQP1* and *AQP5* were almost fully active under NL (each 0.995 ± 0.002) but showed zero activity under Hypo and lower mean activation under HL (0.210 ± 0.409).

**Figure 4.**
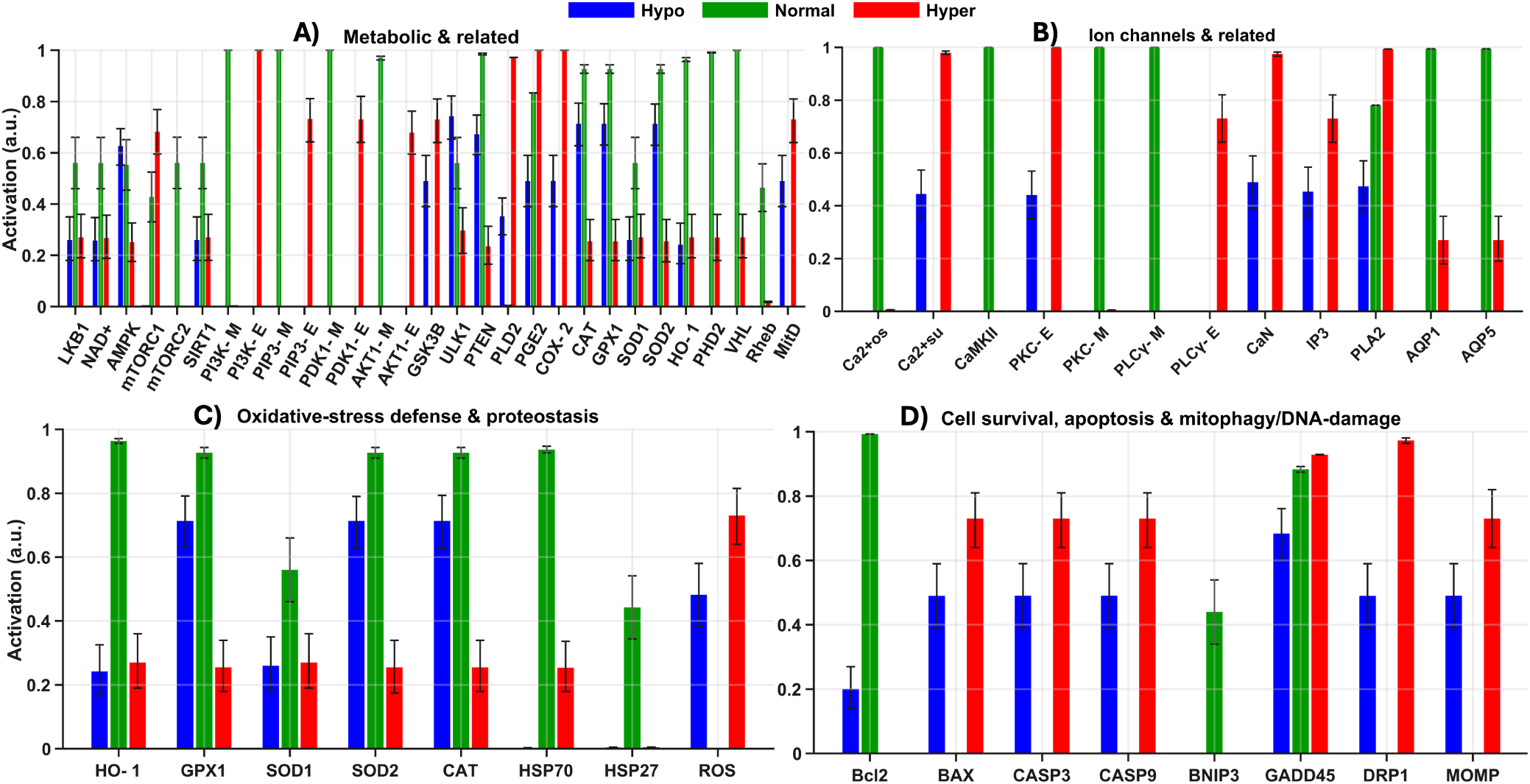
Baseline mean activation under Hypo (blue), NL (green), and HL (red). Bars represent means across 100 replicate simulations per condition; error bars indicate ±1 SD. (A) Metabolic and related signaling nodes; (B) ion channels and calcium-associated nodes; (C) oxidative-stress defense and proteostasis; and (D) cell survival, apoptosis, and mitochondrial-stress markers. Activations are bounded between 0 and 1; symmetric SD whiskers can extend beyond these bounds.

Oxidative-stress and cell-survival readouts differed across the three loading regimes (Fig. 4C–D). Under NL, *HO-1* (0.955 ± 0.038), *GPX1*, *SOD2*, and *CAT* (each 0.908 ± 0.083), and *HSP70*(0.948 ± 0.053) showed high activation, whereas *ROS* was near zero. Under HL, *HO-1* decreased to 0.210 ± 0.409, *GPX1*, *SOD2*, and *CAT* to 0.198 ± 0.386 each, and *HSP70* to 0.198 ± 0.378; *ROS* increased to 0.790 ± 0.409. Under Hypo, the corresponding values were 0.242 ± 0.410 for *HO-1*, 0.750 ± 0.390 for *GPX1*, *SOD2*, and *CAT*, 0.002 ± 0.004 for *HSP70*, and 0.521 ± 0.493 for *ROS*. The survival marker *Bcl2* showed mean activation of 0.992 ± 0.002 under NL, 0.200 ± 0.340 under Hypo, and 0.000 under HL. In contrast, *BAX*, *CASP3*, and *CASP9* were near zero under NL and reached 0.790 ± 0.409 under HL. Under Hypo, *BAX* showed 0.530 ± 0.501, while *CASP3* and *CASP9* each showed 0.530 ± 0.502. All corresponding NL–HL comparisons were significant ( *q <* 0.001).

### 2.3 Model falsification using independent NP mechanobiology experiments

Of the 39 experimental benchmarks, 37 (94.9%) showed RNM-predicted effects concordant with their literature-derived directionality. Agreement was observed for all six benchmarks with positive literature-derived directionality ("+"; 6/6, 100%) and for 31 of the 33 benchmarks with negative directionality ("−"; 31/33, 93.9%). The remaining two benchmarks, HSF1 and CCN2, were contradictory (5.1%), and none was indeterminate. Figure 5 presents the mean steady-state node activations under NL and HL, with SD, for the condition-specific means (panel A), and their qualitative agreement with the literature-derived responses (panel B). In panel B, the upper-left and lower-right triangles represent the RNM-predicted and literature-derived directions, respectively, with green and red indicating relatively higher and lower responses under the corresponding loading condition. The experimental benchmarks and supporting references are provided in Table 2.

**Figure 5.**
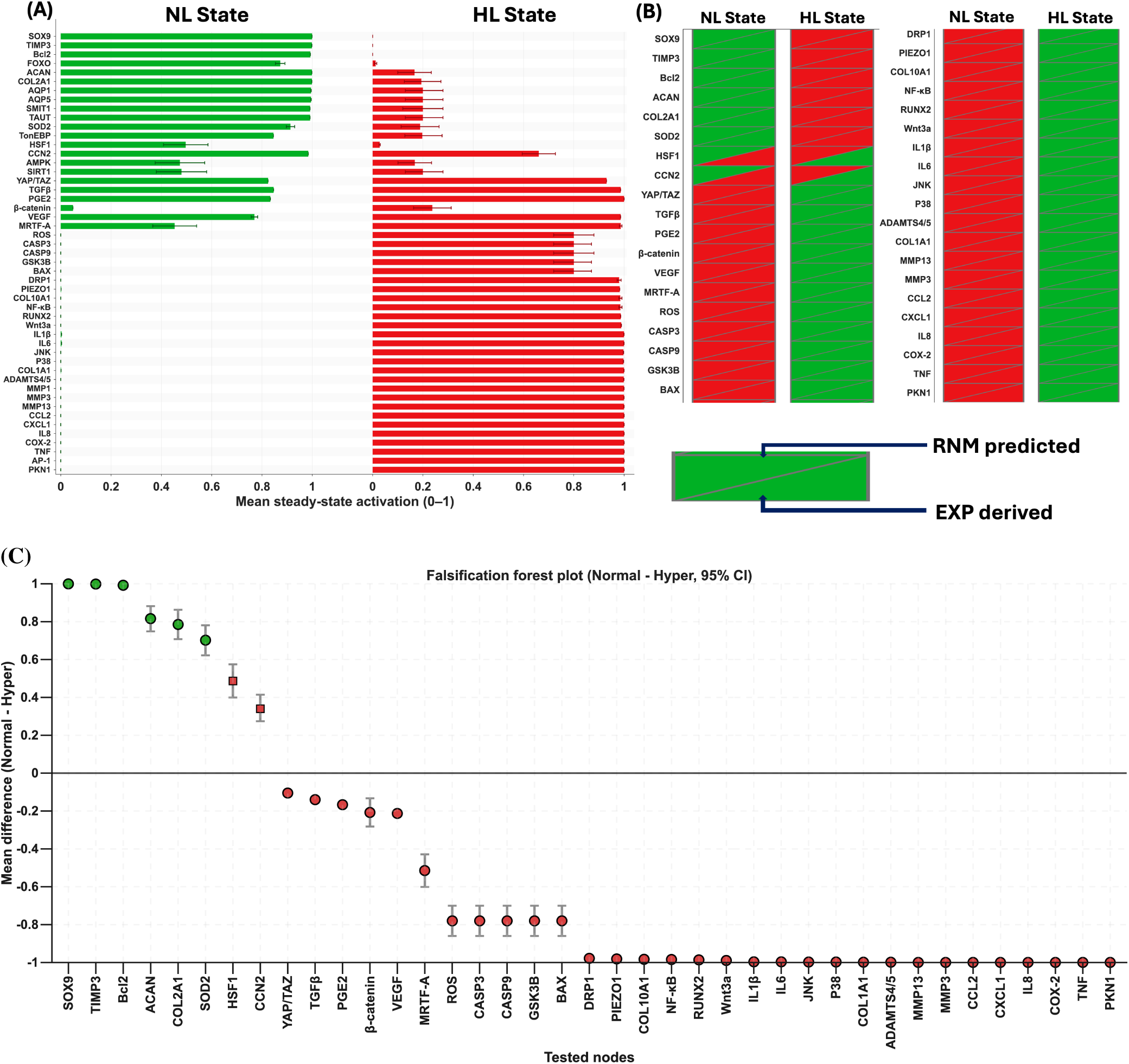
RNM falsification results for 39 experimental benchmarks. (A) Mean steady-state node activation under NL and HL, with standard deviations (SD) across 100 replicate simulations per condition. (B) Directional comparison of RNM predictions and literature-derived responses. (C) Forest plot of RNM-calculated loading effects 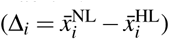 with 95% bootstrap confidence intervals (CIs).

**Table 1.** Representative experimental loading conditions used to guide the interpretation of normal (NL) and hyper/excessive (HL) mechanical loading in the model.

| Study | Experimental model | Experimental load | Interpreted loading |
|---|---|---|---|
| Shah and Chahine, 2018<sup>140</sup> | Bovine NP cells encapsulated in 3D alginate beads | Cyclic hydrostatic pressure of 0–0.1 or 0–0.6 MPa at 0.1 Hz, 2 h/day for 7 days; investigated within the physiological loading range | NL |
| Neidlinger-Wilke et al., 2006<sup>32</sup> | Human and bovine nucleus cells in 3D type-I collagen matrices | Low hydrostatic pressure: 0.25 MPa, 0.1 Hz for 30 min | NL |
| Chooi et al., 2016<sup>141</sup> | Bovine caudal IVD organ culture | Dynamic compression of 0.35 ± 0.25 MPa at 0.2 Hz for 2 h; explicitly applied within the physiological range | NL |
| Shi et al., 2022<sup>115</sup> | Human NP cells in hydrostatic-pressure culture | 3 atm hydrostatic pressure for 2 h; identified as an appropri-ate hydrostatic stimulus with anabolic effects | NL |
| Easson et al., 2023<sup>49</sup> | Mouse functional spine-unit organ culture | Cyclic compression at 10% strain, 1 Hz, 1 h/day for 5 days | NL |
| Yen et al., 2025<sup>142</sup> | Human NP cells in a cyclic-stretch bioreactor | 5% cyclic strain at 0.1 Hz | NL |
| Zhang et al., 2019<sup>27</sup> | Primary human NP cells cultured in BioFlex plates | Each overload cycle consisted of 0.8 MPa at 0.5 Hz for 20 h followed by 0.1 MPa at 0.5 Hz for 4 h; applied for two or five cycles | HL |
| Gawri et al., 2014<sup>29</sup> | Primary human NP and AF cells cul-tured on flexible silicone substrates | High mechanical strain of 20% at 0.001 Hz, applied in 8-h loading bouts separated by 16 h rest | HL |
| Wang et al., 2018<sup>105</sup> | Human NP cells subjected to cyclic mechanical strain | High-amplitude cyclic strain of 19% at 0.01 Hz; prolonged stimulation for up to 72 h | HL |
| Cambria et al., 2021<sup>35</sup> | Bovine NP cells in 3D agarose–collagen hydrogels | Hyperphysiological dynamic compression at 20% strain and 0.5 Hz for 1 h | HL |
| Shi et al., 2022<sup>115</sup> | Human NP cells under hydrostatic pressure | High hydrostatic pressure of 30 atm for 2 h; identified as excessive hydrostatic pressure | HL |
| Wang et al., 2020<sup>88</sup> | Human NP cells encapsulated in 3D GelMA hydrogels | High-magnitude dynamic compression at 20% deformation and 1 Hz for 4 h/day during 5 days of bioreactor culture | HL |
| Li et al., 2017<sup>84</sup> | Rat NP cells seeded in decalcified bone-matrix scaffolds | High-magnitude dynamic compression at 20% deformation and 1 Hz for 6 h/day over 5 days | HL |
| Jia et al., 2024<sup>143</sup> | Primary rat NP cells | Mechanical overloading at 1 MPa and 1 Hz for 1 h | HL |
| Wang et al., 2021<sup>26</sup> | Rabbit lumbar IVD organ culture containing NP and inner AF tissues | Intermittent hydrostatic pressure of 1.0, 3.0, or 5.0 MPa, with 30 s loading and 30 s release cycles for 2 h/day over 30 days; characterized as high-magnitude loading | HL |

**Table 2.**
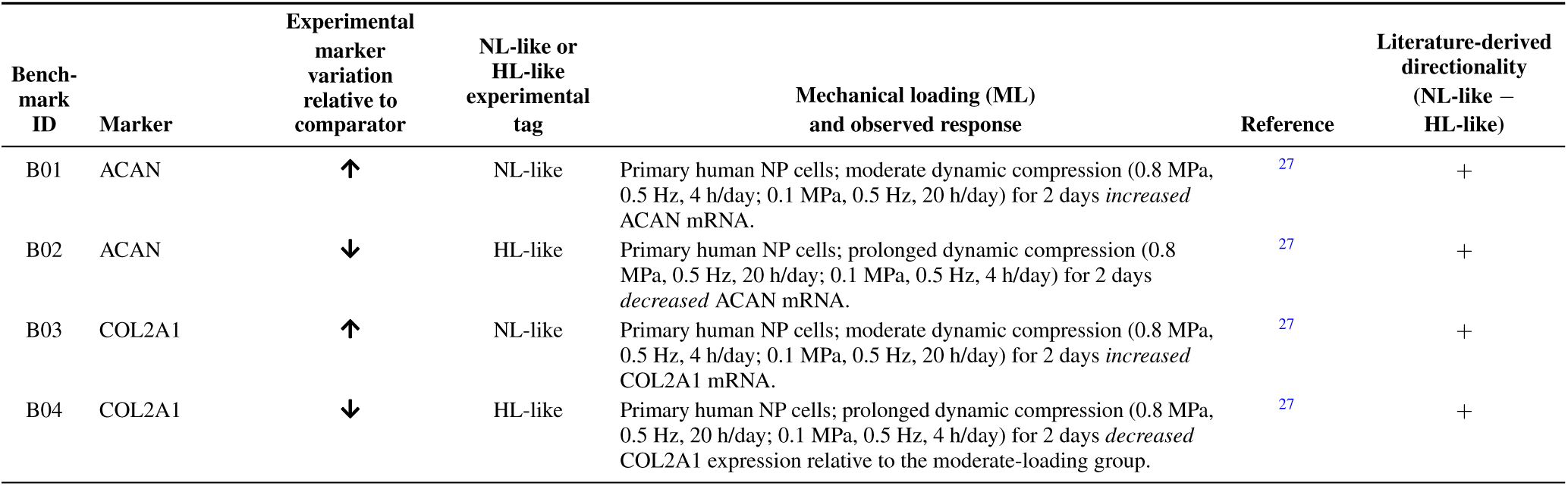
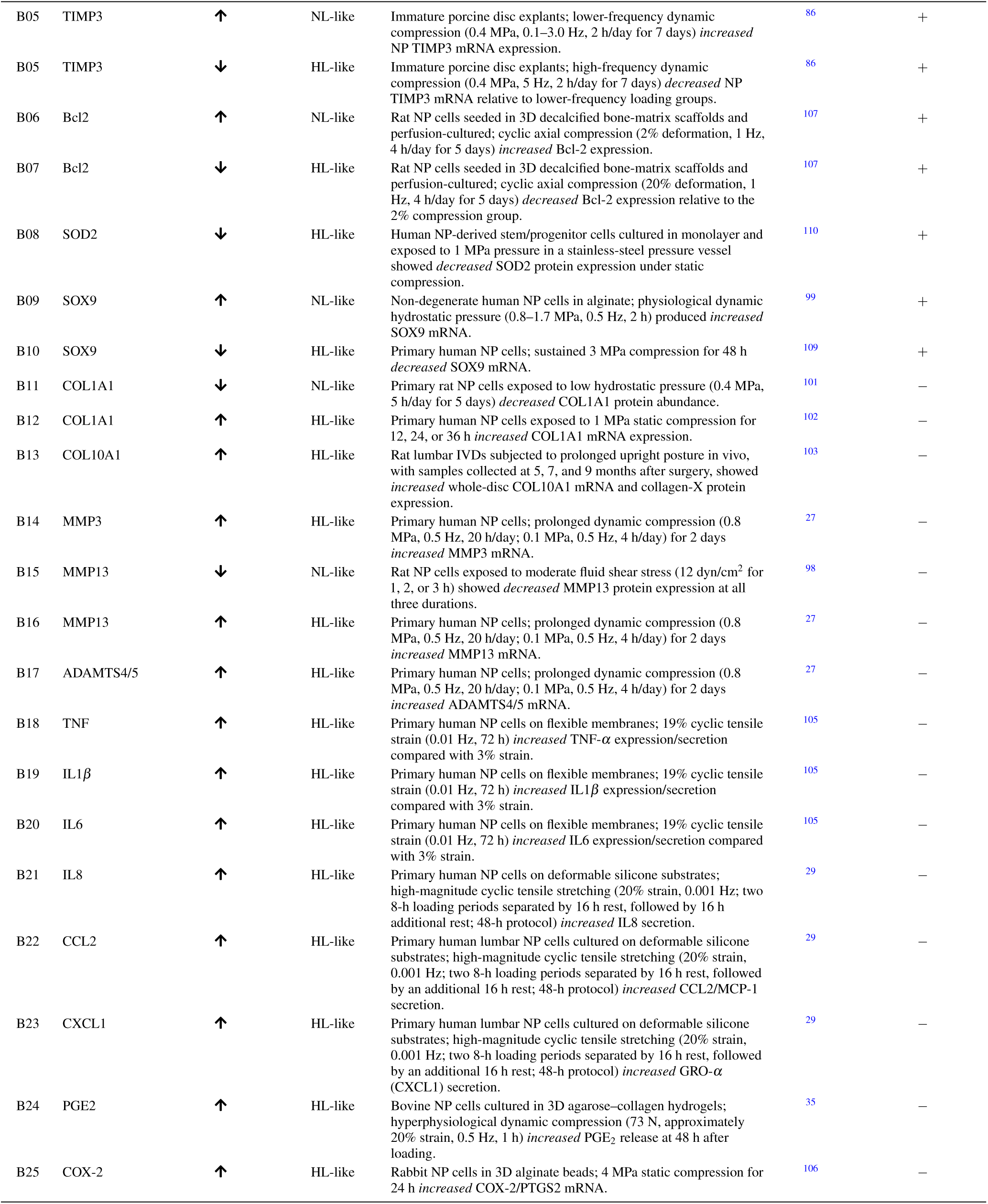
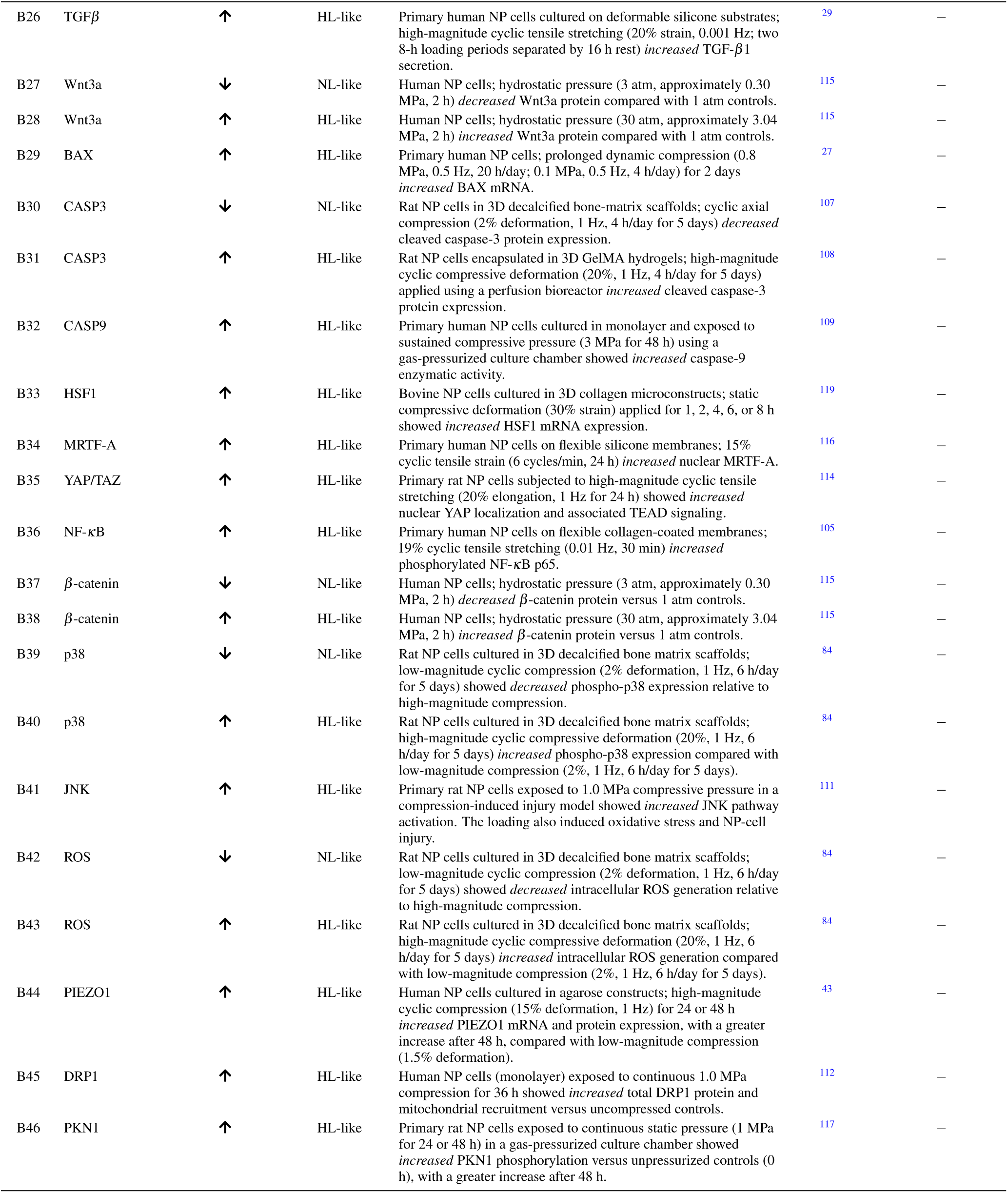
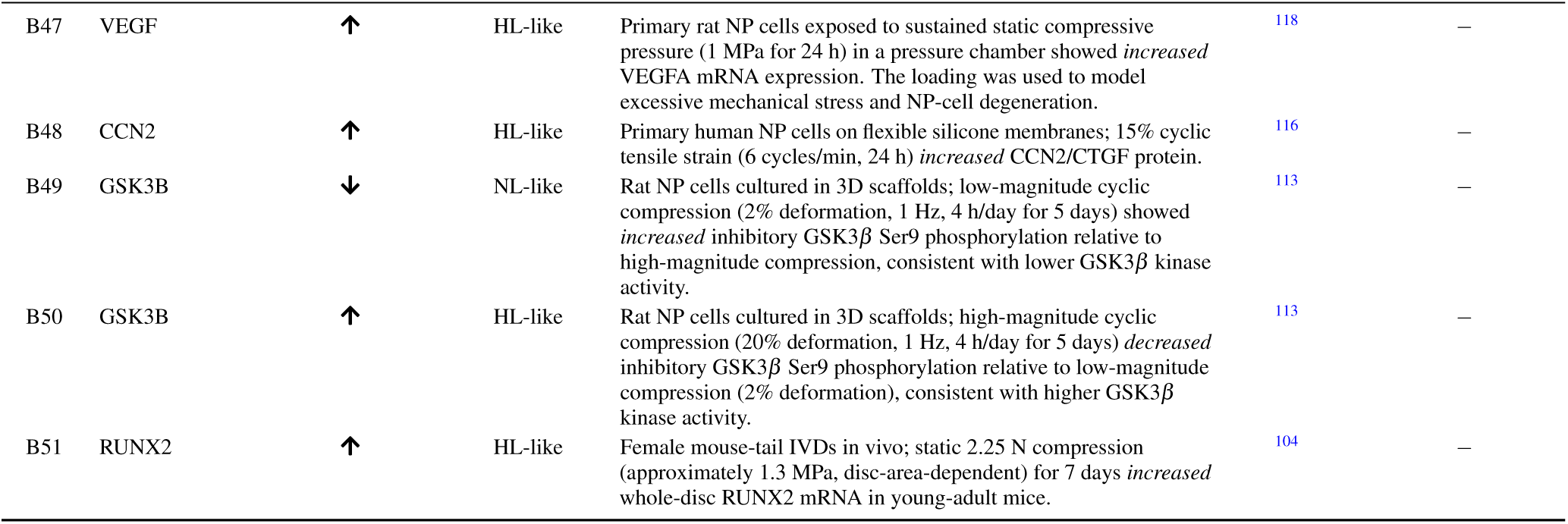
Literature-derived experimental loading-response benchmarks for 39 molecular markers used for independent RNM falsification. Each benchmark entry (B) reports the experimentally observed increase (↑) or decrease (↓) in marker expression or activity under the specified mechanical loading condition, relative to the comparator stated in the corresponding row. Comparators may be unloaded controls or alternative loading conditions. NL-like and HL-like denote physiological-like and excessive-like experimental loading regimes, respectively. The final column specifies the literature-derived marker-level directionality used for falsification: + indicates higher expression or activity under NL-like loading relative to HL-like loading, whereas − indicates higher expression or activity under HL-like loading relative to NL-like loading.

| Bench-mark ID | Marker | Experimental marker variation relative to comparator | NL-like or HL-like experimental tag | Mechanical loading (ML) and observed response | Reference | Literature-derived directionality (NL-like − HL-like) |
|---|---|---|---|---|---|---|
| B01 | ACAN | ↑ | NL-like | Primary human NP cells; moderate dynamic compression (0.8 MPa, 0.5 Hz, 4 h/day; 0.1 MPa, 0.5 Hz, 20 h/day) for 2 days *increased* ACAN mRNA. | <sup>27</sup> | + |
| B02 | ACAN | ↓ | HL-like | Primary human NP cells; prolonged dynamic compression (0.8 MPa, 0.5 Hz, 20 h/day; 0.1 MPa, 0.5 Hz, 4 h/day) for 2 days *decreased* ACAN mRNA. | <sup>27</sup> | + |
| B03 | COL2A1 | ↑ | NL-like | Primary human NP cells; moderate dynamic compression (0.8 MPa, 0.5 Hz, 4 h/day; 0.1 MPa, 0.5 Hz, 20 h/day) for 2 days *increased* COL2A1 mRNA. | <sup>27</sup> | + |
| B04 | COL2A1 | ↓ | HL-like | Primary human NP cells; prolonged dynamic compression (0.8 MPa, 0.5 Hz, 20 h/day; 0.1 MPa, 0.5 Hz, 4 h/day) for 2 days *decreased* COL2A1 expression relative to the moderate-loading group. | <sup>27</sup> | + |
| B05 | TIMP3 | ↑ | NL-like | Immature porcine disc explants; lower-frequency dynamic compression (0.4 MPa, 0.1–3.0 Hz, 2 h/day for 7 days) *increased* NP TIMP3 mRNA expression. | <sup>86</sup> | + |
| B05 | TIMP3 | ↓ | HL-like | Immature porcine disc explants; high-frequency dynamic compression (0.4 MPa, 5 Hz, 2 h/day for 7 days) *decreased* NP TIMP3 mRNA relative to lower-frequency loading groups. | <sup>86</sup> | + |
| B06 | Bcl2 | ↑ | NL-like | Rat NP cells seeded in 3D decalcified bone-matrix scaffolds and perfusion-cultured; cyclic axial compression (2% deformation, 1 Hz, 4 h/day for 5 days) *increased* Bcl-2 expression. | <sup>107</sup> | + |
| B07 | Bcl2 | ↓ | HL-like | Rat NP cells seeded in 3D decalcified bone-matrix scaffolds and perfusion-cultured; cyclic axial compression (20% deformation, 1 Hz, 4 h/day for 5 days) *decreased* Bcl-2 expression relative to the 2% compression group. | <sup>107</sup> | + |
| B08 | SOD2 | ↓ | HL-like | Human NP-derived stem/progenitor cells cultured in monolayer and exposed to 1 MPa pressure in a stainless-steel pressure vessel showed *decreased* SOD2 protein expression under static compression. | <sup>110</sup> | + |
| B09 | SOX9 | ↑ | NL-like | Non-degenerate human NP cells in alginate; physiological dynamic hydrostatic pressure (0.8–1.7 MPa, 0.5 Hz, 2 h) produced *increased* SOX9 mRNA. | <sup>99</sup> | + |
| B10 | SOX9 | ↓ | HL-like | Primary human NP cells; sustained 3 MPa compression for 48 h *decreased* SOX9 mRNA. | <sup>109</sup> | + |
| B11 | COL1A1 | ↓ | NL-like | Primary rat NP cells exposed to low hydrostatic pressure (0.4 MPa, 5 h/day for 5 days) *decreased* COL1A1 protein abundance. | <sup>101</sup> | − |
| B12 | COL1A1 | ↑ | HL-like | Primary human NP cells exposed to 1 MPa static compression for 12, 24, or 36 h *increased* COL1A1 mRNA expression. | <sup>102</sup> | − |
| B13 | COL10A1 | ↑ | HL-like | Rat lumbar IVDs subjected to prolonged upright posture in vivo, with samples collected at 5, 7, and 9 months after surgery, showed *increased* whole-disc COL10A1 mRNA and collagen-X protein expression. | <sup>103</sup> | − |
| B14 | MMP3 | ↑ | HL-like | Primary human NP cells; prolonged dynamic compression (0.8 MPa, 0.5 Hz, 20 h/day; 0.1 MPa, 0.5 Hz, 4 h/day) for 2 days *increased* MMP3 mRNA. | <sup>27</sup> | − |
| B15 | MMP13 | ↓ | NL-like | Rat NP cells exposed to moderate fluid shear stress (12 dyn/cm<sup>2</sup> for 1, 2, or 3 h) showed *decreased* MMP13 protein expression at all three durations. | <sup>98</sup> | − |
| B16 | MMP13 | ↑ | HL-like | Primary human NP cells; prolonged dynamic compression (0.8 MPa, 0.5 Hz, 20 h/day; 0.1 MPa, 0.5 Hz, 4 h/day) for 2 days *increased* MMP13 mRNA. | <sup>27</sup> | − |
| B17 | ADAMTS4/5 | ↑ | HL-like | Primary human NP cells; prolonged dynamic compression (0.8 MPa, 0.5 Hz, 20 h/day; 0.1 MPa, 0.5 Hz, 4 h/day) for 2 days *increased* ADAMTS4/5 mRNA. | <sup>27</sup> | − |
| B18 | TNF | ↑ | HL-like | Primary human NP cells on flexible membranes; 19% cyclic tensile strain (0.01 Hz, 72 h) *increased* TNF-α expression/secretion compared with 3% strain. | <sup>105</sup> | − |
| B19 | IL1β | ↑ | HL-like | Primary human NP cells on flexible membranes; 19% cyclic tensile strain (0.01 Hz, 72 h) *increased* IL1β expression/secretion compared with 3% strain. | <sup>105</sup> | − |
| B20 | IL6 | ↑ | HL-like | Primary human NP cells on flexible membranes; 19% cyclic tensile strain (0.01 Hz, 72 h) *increased* IL6 expression/secretion compared with 3% strain. | <sup>105</sup> | − |
| B21 | IL8 | ↑ | HL-like | Primary human NP cells on deformable silicone substrates; high-magnitude cyclic tensile stretching (20% strain, 0.001 Hz; two 8-h loading periods separated by 16 h rest, followed by 16 h additional rest; 48-h protocol) *increased* IL8 secretion. | <sup>29</sup> | − |
| B22 | CCL2 | ↑ | HL-like | Primary human lumbar NP cells cultured on deformable silicone substrates; high-magnitude cyclic tensile stretching (20% strain, 0.001 Hz; two 8-h loading periods separated by 16 h rest, followed by an additional 16 h rest; 48-h protocol) *increased* CCL2/MCP-1 secretion. | <sup>29</sup> | − |
| B23 | CXCL1 | ↑ | HL-like | Primary human lumbar NP cells cultured on deformable silicone substrates; high-magnitude cyclic tensile stretching (20% strain, 0.001 Hz; two 8-h loading periods separated by 16 h rest, followed by an additional 16 h rest; 48-h protocol) *increased* GRO-α (CXCL1) secretion. | <sup>29</sup> | − |
| B24 | PGE2 | ↑ | HL-like | Bovine NP cells cultured in 3D agarose–collagen hydrogels; hyperphysiological dynamic compression (73 N, approximately 20% strain, 0.5 Hz, 1 h) *increased* PGE<sub>2</sub> release at 48 h after loading. | <sup>35</sup> | − |
| B25 | COX-2 | ↑ | HL-like | Rabbit NP cells in 3D alginate beads; 4 MPa static compression for 24 h *increased* COX-2/PTGS2 mRNA. | <sup>106</sup> | − |
| B26 | TGFβ | ↑ | HL-like | Primary human NP cells cultured on deformable silicone substrates; high-magnitude cyclic tensile stretching (20% strain, 0.001 Hz; two 8-h loading periods separated by 16 h rest) *increased* TGF-β1 secretion. | <sup>29</sup> | − |
| B27 | Wnt3a | ↓ | NL-like | Human NP cells; hydrostatic pressure (3 atm, approximately 0.30 MPa, 2 h) *decreased* Wnt3a protein compared with 1 atm controls. | <sup>115</sup> | − |
| B28 | Wnt3a | ↑ | HL-like | Human NP cells; hydrostatic pressure (30 atm, approximately 3.04 MPa, 2 h) *increased* Wnt3a protein compared with 1 atm controls. | <sup>115</sup> | − |
| B29 | BAX | ↑ | HL-like | Primary human NP cells; prolonged dynamic compression (0.8 MPa, 0.5 Hz, 20 h/day; 0.1 MPa, 0.5 Hz, 4 h/day) for 2 days *increased* BAX mRNA. | <sup>27</sup> | − |
| B30 | CASP3 | ↓ | NL-like | Rat NP cells in 3D decalcified bone-matrix scaffolds; cyclic axial compression (2% deformation, 1 Hz, 4 h/day for 5 days) *decreased* cleaved caspase-3 protein expression. | <sup>107</sup> | − |
| B31 | CASP3 | ↑ | HL-like | Rat NP cells encapsulated in 3D GelMA hydrogels; high-magnitude cyclic compressive deformation (20%, 1 Hz, 4 h/day for 5 days) applied using a perfusion bioreactor *increased* cleaved caspase-3 protein expression. | <sup>108</sup> | − |
| B32 | CASP9 | ↑ | HL-like | Primary human NP cells cultured in monolayer and exposed to sustained compressive pressure (3 MPa for 48 h) using a gas-pressurized culture chamber showed *increased* caspase-9 enzymatic activity. | <sup>109</sup> | − |
| B33 | HSF1 | ↑ | HL-like | Bovine NP cells cultured in 3D collagen microconstructs; static compressive deformation (30% strain) applied for 1, 2, 4, 6, or 8 h showed *increased* HSF1 mRNA expression. | <sup>119</sup> | − |
| B34 | MRTF-A | ↑ | HL-like | Primary human NP cells on flexible silicone membranes; 15% cyclic tensile strain (6 cycles/min, 24 h) *increased* nuclear MRTF-A. | <sup>116</sup> | − |
| B35 | YAP/TAZ | ↑ | HL-like | Primary rat NP cells subjected to high-magnitude cyclic tensile stretching (20% elongation, 1 Hz for 24 h) showed *increased* nuclear YAP localization and associated TEAD signaling. | <sup>114</sup> | − |
| B36 | NF-κB | ↑ | HL-like | Primary human NP cells on flexible collagen-coated membranes; 19% cyclic tensile stretching (0.01 Hz, 30 min) *increased* phosphorylated NF-κB p65. | <sup>105</sup> | − |
| B37 | β-catenin | ↓ | NL-like | Human NP cells; hydrostatic pressure (3 atm, approximately 0.30 MPa, 2 h) *decreased* β-catenin protein versus 1 atm controls. | <sup>115</sup> | − |
| B38 | β-catenin | ↑ | HL-like | Human NP cells; hydrostatic pressure (30 atm, approximately 3.04 MPa, 2 h) *increased* β-catenin protein versus 1 atm controls. | <sup>115</sup> | − |
| B39 | p38 | ↓ | NL-like | Rat NP cells cultured in 3D decalcified bone matrix scaffolds; low-magnitude cyclic compression (2% deformation, 1 Hz, 6 h/day for 5 days) showed *decreased* phospho-p38 expression relative to high-magnitude compression. | <sup>84</sup> | − |
| B40 | p38 | ↑ | HL-like | Rat NP cells cultured in 3D decalcified bone matrix scaffolds; high-magnitude cyclic compressive deformation (20%, 1 Hz, 6 h/day for 5 days) *increased* phospho-p38 expression compared with low-magnitude compression (2%, 1 Hz, 6 h/day for 5 days). | <sup>84</sup> | − |
| B41 | JNK | ↑ | HL-like | Primary rat NP cells exposed to 1.0 MPa compressive pressure in a compression-induced injury model showed *increased* JNK pathway activation. The loading also induced oxidative stress and NP-cell injury. | <sup>111</sup> | − |
| B42 | ROS | ↓ | NL-like | Rat NP cells cultured in 3D decalcified bone matrix scaffolds; low-magnitude cyclic compression (2% deformation, 1 Hz, 6 h/day for 5 days) showed *decreased* intracellular ROS generation relative to high-magnitude compression. | <sup>84</sup> | − |
| B43 | ROS | ↑ | HL-like | Rat NP cells cultured in 3D decalcified bone matrix scaffolds; high-magnitude cyclic compressive deformation (20%, 1 Hz, 6 h/day for 5 days) *increased* intracellular ROS generation compared with low-magnitude compression (2%, 1 Hz, 6 h/day for 5 days). | <sup>84</sup> | − |
| B44 | PIEZO1 | ↑ | HL-like | Human NP cells cultured in agarose constructs; high-magnitude cyclic compression (15% deformation, 1 Hz) for 24 or 48 h *increased* PIEZO1 mRNA and protein expression, with a greater increase after 48 h, compared with low-magnitude compression (1.5% deformation). | <sup>43</sup> | − |
| B45 | DRP1 | ↑ | HL-like | Human NP cells (monolayer) exposed to continuous 1.0 MPa compression for 36 h showed *increased* total DRP1 protein and mitochondrial recruitment versus uncompressed controls. | <sup>112</sup> | − |
| B46 | PKN1 | ↑ | HL-like | Primary rat NP cells exposed to continuous static pressure (1 MPa for 24 or 48 h) in a gas-pressurized culture chamber showed *increased* PKN1 phosphorylation versus unpressurized controls (0 h), with a greater increase after 48 h. | <sup>117</sup> | − |
| B47 | VEGF | ↑ | HL-like | Primary rat NP cells exposed to sustained static compressive pressure (1 MPa for 24 h) in a pressure chamber showed *increased* VEGFA mRNA expression. The loading was used to model excessive mechanical stress and NP-cell degeneration. | <sup>118</sup> | − |
| B48 | CCN2 | ↑ | HL-like | Primary human NP cells on flexible silicone membranes; 15% cyclic tensile strain (6 cycles/min, 24 h) *increased* CCN2/CTGF protein. | <sup>116</sup> | − |
| B49 | GSK3B | ↓ | NL-like | Rat NP cells cultured in 3D scaffolds; low-magnitude cyclic compression (2% deformation, 1 Hz, 4 h/day for 5 days) showed *increased* inhibitory GSK3β Ser9 phosphorylation relative to high-magnitude compression, consistent with lower GSK3β kinase activity. | <sup>113</sup> | − |
| B50 | GSK3B | ↑ | HL-like | Rat NP cells cultured in 3D scaffolds; high-magnitude cyclic compression (20% deformation, 1 Hz, 4 h/day for 5 days) *decreased* inhibitory GSK3β Ser9 phosphorylation relative to low-magnitude compression (2% deformation), consistent with higher GSK3β kinase activity. | <sup>113</sup> | − |
| B51 | RUNX2 | ↑ | HL-like | Female mouse-tail IVDs in vivo; static 2.25 N compression (approximately 1.3 MPa, disc-area-dependent) for 7 days *increased* whole-disc RUNX2 mRNA in young-adult mice. | <sup>104</sup> | − |

The bootstrap forest plot (Fig. 5C) presents the model-calculated loading effects 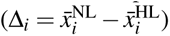 and their 95% percentile bootstrap CIs. Positive and negative values indicate higher simulated activation under NL and HL, respectively. For the 6 benchmarks with "+" literature-derived response, the effects (95% CIs) were SOX9 (+0.9993 [+0.9993, +0.9994]), TIMP3 (+0.9983 [+0.9980, +0.9985]), Bcl2 (+0.9925 [+0.9922, +0.9928]), ACAN (+0.8154 [+0.7488, +0.8820]), COL2A1 (+0.7851 [+0.7075, +0.8627]), and SOD2 (+0.7025 [+0.6221, +0.7807]). Among the benchmarks with "−" literature- derived response, representative concordant effects included COL10A1 (−0.9835 [−0.9895, −0.9776]), MMP3 (−0.99946 [−0.99950, −0.99942]), IL1*β* (−0.9963 [−0.9968, −0.9957]), NF-*κ*B (−0.9838 [−0.9897, −0.9778]), VEGF (−0.2132 [−0.2260, −0.2004]), MRTF-A (−0.5145 [−0.6009, −0.4287]), *β* -catenin (−0.2074 [−0.2821, −0.1328]), BAX (−0.7800 [−0.8600, −0.7000]), DRP1 (−0.9778 [−0.9859, −0.9698]), GSK3B (−0.7800 [−0.8600, −0.7000]), and ROS (−0.7800 [−0.8600, −0.7000]). The two contradictory benchmarks showed positive effects: HSF1 (+0.4872 [+0.3995, +0.5746]) and CCN2 (+0.3401 [+0.2741, +0.4144]). In the forest plot, green and red markers distinguish benchmarks with upward and downward literature-derived responses, respectively, while square markers identify the contradictory predictions. The complete numerical results and individual classifications are reported in Supplementary Table S2.

### 2.4 Hyper-to-Normal rescue screening via node clamping

The 48 single- and dual-node clamping strategies produced different degrees of HL-to-NL profile recovery across the eight functional modules. The highest module-specific scores ranged from 10.4% for ion channels and related nodes to 85.9% for ECM anabolism and phenotype markers (Figs. 6–7; Supplementary Table S3). Here, recovery denotes the percentage reduction in the Euclidean distance from the intervention profile to the NL reference, normalized to the original HL-to-NL distance within each module.

**Figure 6.**
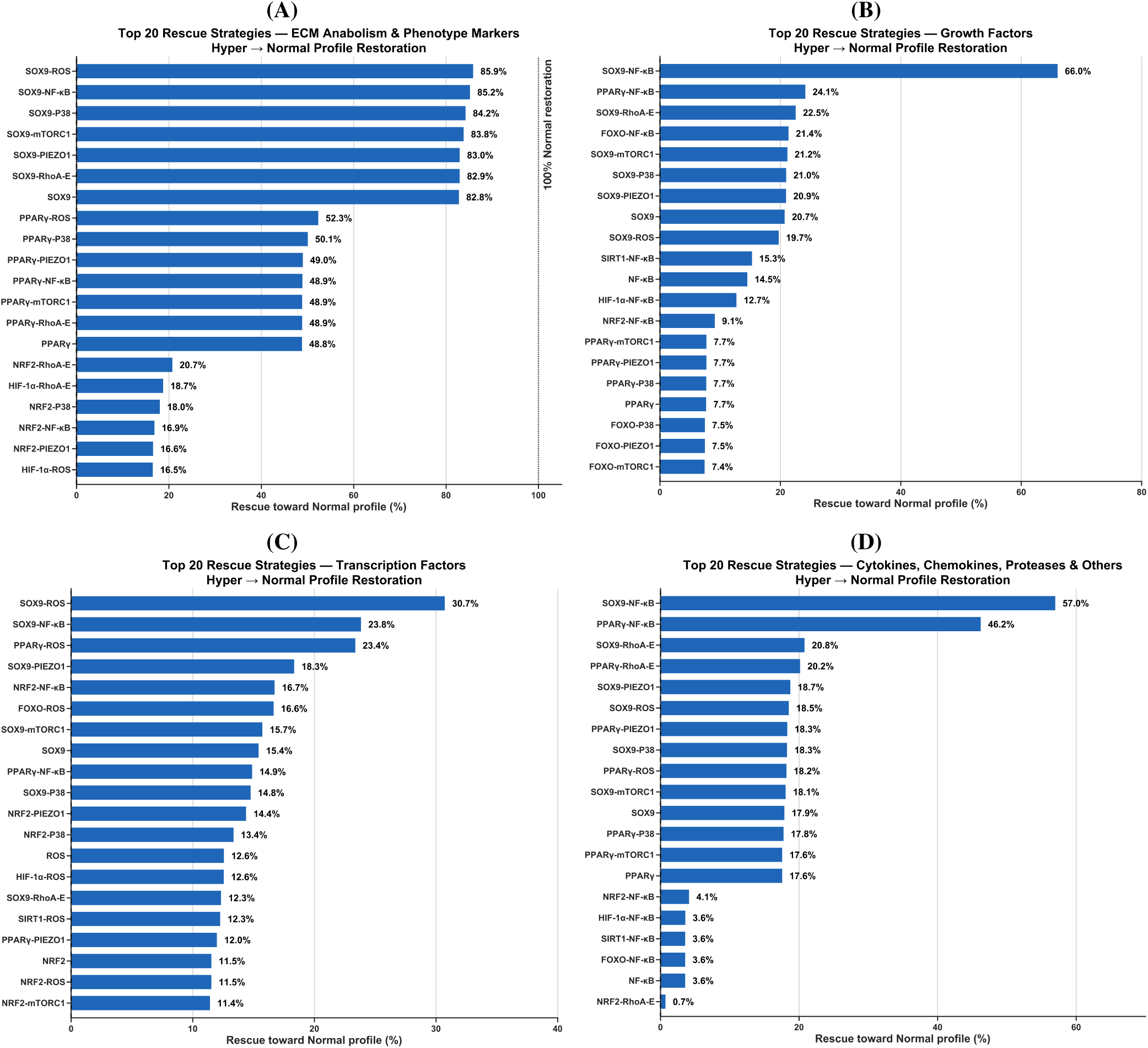
Top 20 node-clamping strategies ranked by HL-to-NL profile recovery in four modules: (A) ECM anabolism and phenotype markers, (B) growth factors, (C) transcription factors, and (D) cytokines, chemokines, proteases, and related mediators. Each percentage is the reduction in the module-specific Euclidean distance to the NL reference relative to the unperturbed HL-to-NL distance.

**Figure 7.**
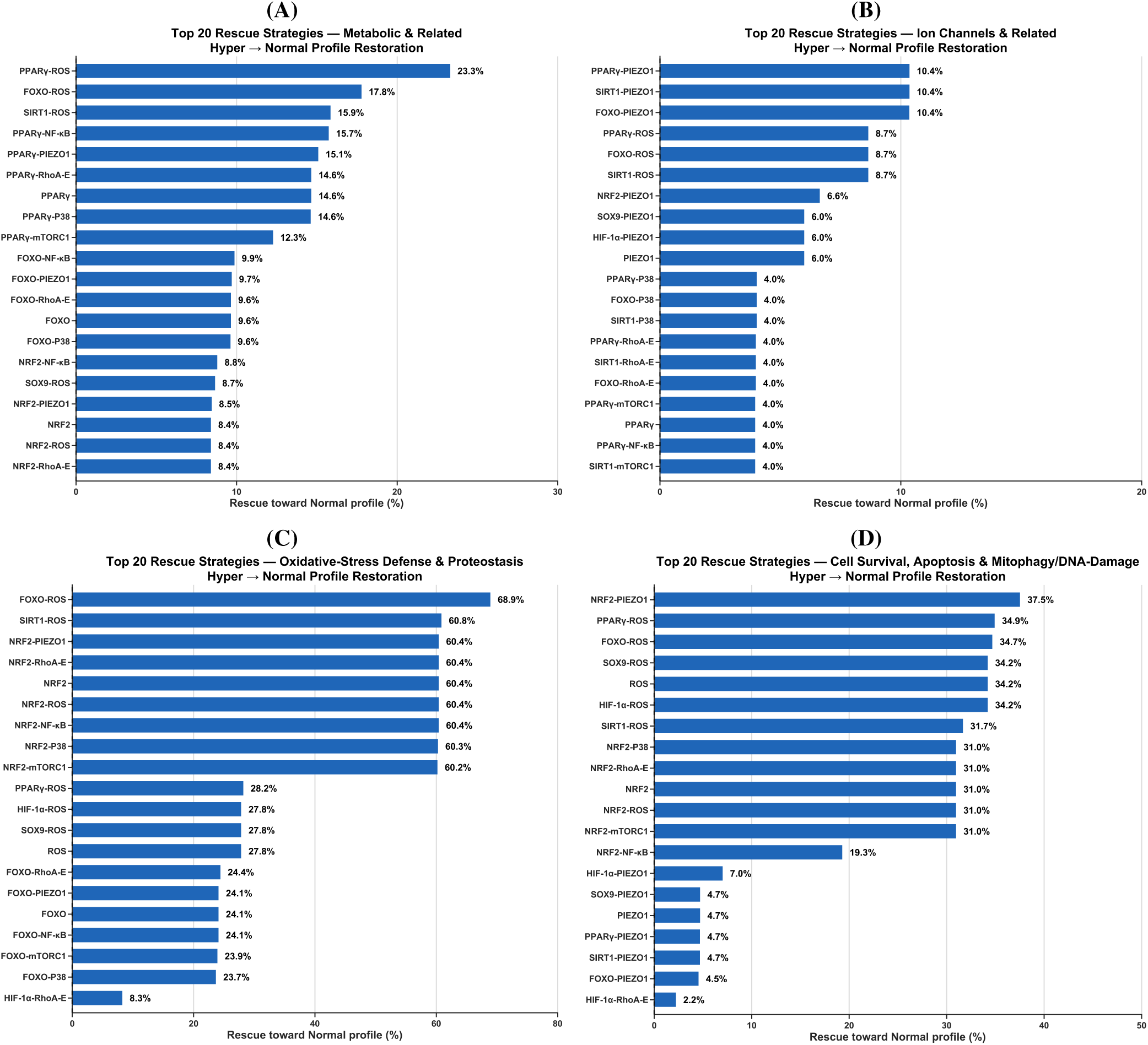
Top 20 node-clamping strategies ranked by HL-to-NL profile recovery in four modules: (A) metabolic and related nodes, (B) ion channels and related nodes, (C) oxidative-stress defense and proteostasis, and (D) cell survival, apoptosis, mitophagy, and DNA-damage responses. Recovery percentages are defined relative to the original module-specific HL-to-NL Euclidean distance.

The ECM module showed the largest recovery scores (Fig. 6A). SOX9 activation combined with ROS suppression ranked first (85.9%), followed by SOX9 activation paired with NF-*κ*B (85.2%), p38 (84.2%), or mTORC1 suppression (83.8%). Combinations targeting PIEZO1 and RhoA-E achieved 83.0% and 82.9%, respectively. SOX9 activation alone yielded 82.8%, only 3.1 percentage points below the leading dual intervention. By comparison, PPAR*γ* activation alone achieved 48.8%, increasing to 52.3% with ROS suppression. In the growth-factor module (Fig. 6B), SOX9 activation with NF-*κ*B suppression achieved 66.0%, compared with 24.1% for PPAR*γ*–NF-*κ*B, 22.5% for SOX9–RhoA-E, and 20.7% for SOX9 alone. The transcription-factor and cytokine/protease modules showed different leading combinations (Fig. 6C,D). In transcription factors, SOX9–ROS achieved 30.7% recovery, followed by SOX9–NF-*κ*B (23.8%) and PPAR*γ*–ROS (23.4%); the respective single-node scores for SOX9 and ROS were 15.4% and 12.6%. For cytokines, chemokines, proteases, and related mediators, SOX9–NF-*κ*B ranked first (57.0%), followed by PPAR*γ*–NF-*κ*B (46.2%). The next-ranked combinations, SOX9–RhoA-E and PPAR*γ*–RhoA-E, achieved 20.8% and 20.2%, whereas SOX9, PPAR*γ*, and NF-*κ*B single-node interventions achieved 17.9%, 17.6%, and 3.6%, respectively.

The metabolic module showed lower recovery, led by PPAR*γ*–ROS (23.3%), FOXO–ROS (17.8%), and SIRT1–ROS (15.9%) (Fig. 7A). PPAR*γ* activation alone achieved 14.6%. In the ion-channel module, PPAR*γ*–PIEZO1, SIRT1–PIEZO1, and FOXO–PIEZO1 shared the highest reported score (10.4%), compared with 6.0% for PIEZO1 suppression alone; the corresponding ROS-containing combinations each achieved 8.7% (Fig. 7B). Oxidative-stress defense and proteostasis showed 68.9% recovery with FOXO activation and ROS suppression, followed by SIRT1–ROS (60.8%) (Fig. 7C). NRF2 activation alone achieved 60.4%, and several NRF2-containing pairs clustered at 60.2–60.4%. The corresponding FOXO-only and ROS-only scores were 24.1% and 27.8%. For cell survival, apoptosis, mitophagy, and DNA-damage nodes (Fig. 7D), NRF2–PIEZO1 ranked first (37.5%), compared with 31.0% for NRF2 activation alone. PPAR*γ*–ROS (34.9%) and FOXO–ROS (34.7%) followed, while ROS suppression alone achieved 34.2%.

Node-resolved responses for the two leading strategies in each module are reported in Supplementary Figs. S2 and S3. In the ECM module, SOX9 activation with ROS suppression increased the mean activation of ACAN, COL2A1, and TIMP3 from 0.216, 0.252, and less than 0.001 under HL to 0.986, 0.994, and 0.716, respectively. In the cytokine/protease module, SOX9 activation with NF-*κ*B suppression reduced IL8, CCL2, CXCL1, and CXCL3 from approximately 1.000 to 0.008 and MMP1, MMP3, and MMP13 from approximately 0.999 to below 0.001. Across the predefined eight-marker anabolic and ten-marker catabolic panels, mean activations were 0.985 and 0.001 under NL and 0.059 and 0.973 under HL. The corresponding means were 0.589 and 0.653 for SOX9–ROS and 0.718 and 0.371 for SOX9–NF-*κ*B (Supplementary Table S4).

## 3 Discussion

### 3.1 Network architecture and structural features

A central contribution of this study is the consolidation of dispersed NP mechanobiology into a single signed, directed regulatory network that connects mechanosensory entry points to transcriptional, metabolic, inflammatory, and matrix-maintenance outputs (Fig. 1). The resulting architecture is sparse (network density = 0.033; average node degree = 4.803), yet exhibits relatively short directed paths (characteristic path length = 3.363; network diameter = 8.00) and non-random local connectivity (low clustering coefficient = 0.103, high network heterogeneity = 0.906, low centralization =0.154), consistent with a signalling system that may rapidly integrate mechanical cues into coordinated cellular decisions within a constrained microenvironment^72,75–77^. These topological properties provide a structural basis that supports the emergence of distinct load-dependent states and a limited set of effective intervention points observed in the model, as further discussed below.

Centrality analysis suggests a layered control organization. First, a small set of *broadcasting regulators* (high signed out-degree) can influence many downstream targets and thereby consolidate phenotype-level programs. The high outgoing influence of the NF-*κ*B protein complex node and AP-1 transcription factor node (Fig. 2A) is consistent with the respective roles of these cell signalling components as inflammatory amplifiers and broad transcriptional regulators of catabolic remodelling^50,78^. In turn, SOX9 and PPAR*γ* appear as prominent outgoing regulators aligned with the stabilization of a normal NP phenotype and suppression of inflammatory and catabolic responses^79,80^. Accordingly, they will emerge as key factors for the simulated cell rescues in HL environments, which will be discussed later. Second, *routing hubs* (high betweenness), the nodes of the AMPK–mTOR and PI3K–AKT axes (Fig. 2B), appeared topologically well positioned to gate information flow between mechanosensory modules and downstream transcriptional outputs. This is consistent with experimental evidence that metabolic signalling couples mechanical cues with energetic and redox constraints, thereby shaping survival and stress adaptation^81,82^.

Third, *globally accessible nodes* (high harmonic closeness), such as ROS, ERK, Ca^2+^-related mediators, and FAK-associated modules (Fig. 2C), are topologically positioned to influence multiple subsystems with minimal path length. This aligns with established roles of integrin–FAK signalling, mechanosensitive Ca^2+^ responses, MAPK activation, and ROS-dependent stress signalling in NP mechanoadaptation and degeneration^23,35,38,83,84^, and it allowed to identify several related nodes as important complementary targets in the rescue analyses.

Overall, the layered organization of the network achieves through knowledge curation and projection, is consistent with a hierarchical interpretation of NP mechanotransduction in which mechanical inputs are sensed through adhesion- and ion- channel–mediated pathways, routed through metabolic and stress-response modules, and consolidated by a limited set of transcriptional and redox regulators. Nodes positioned near the top of this hierarchy (e.g., adhesion-associated signalling modules) or those stabilizing global stress-buffering programs (e.g., NRF2) were therefore expected to exert broad system-level influence when perturbed^85^. This structural perspective provides a promising framework for mechanistic interpretations of both the load-dependent network states and the sensitivity of the system to targeted regulatory interventions, as further developed through this discussion.

### 3.2 Baseline mechanotransductive phenotypes under Hypo, Normal, and Hyper loading

The baseline profiles describe three regulatory configurations of the NP MT network (Figs. 3–4). NL favors an ECM-maintaining phenotype supported by mechanosensitive, metabolic, and stress-adaptive regulation. HL combines altered matrix regulation with inflammatory and mitochondrial stress, while Hypo couples insufficient mechanical stimulation to a distinct intracellular stress response. These configurations are consistent with the loading-dependent nature of NP matrix homeostasis^15,22^.

The central feature of the NL phenotype is its coordinated matrix regulation. The RNM maintained high ACAN (0.999), COL2A1 (0.998), SOX9 (0.999), and TIMP3 (0.998) activation, alongside low COL1A1 (0.001) and matrix-protease activity. This combination represents preservation of an aggrecan- and type-II-collagen-associated NP phenotype with suppression of excessive matrix breakdown. Li et al. examined immature porcine discs under dynamic compression and reported higher aggrecan and collagen-II expression at 0.1–0.8 MPa, with greater TIMP3 expression at 0.1–0.4 MPa; at 1.3 MPa, matrix-gene expression declined as MMP3 and ADAMTS4 increased^86^. The model’s NL-associated matrix pattern therefore corresponds to the experimental relationship between suitable mechanical stimulation and balanced matrix synthesis and degradation. Gao et al. further showed that periodic loading increased aggrecan and collagen-II expression in rat NP cells, while inhibition of integrin-*α*1 or PLC*γ*1 diminished these responses^28^. This links the predicted ECM phenotype to active cell–matrix mechanotransduction.

The pathways accompanying the NL matrix phenotype provide a plausible basis for its maintenance. The RNM showed near-maximal PLC*γ*-M and CaMKII activation, with AKT1-M at 0.964 ± 0.035, Bcl2 at 0.992 ± 0.002, and strong antioxidant activity. Xu et al. found that dynamic compression maintained aggrecan and collagen-II production in three-dimensional rat NP-cell cultures through N-cadherin-associated PI3K–AKT signaling; N-cadherin knockdown or PI3K inhibition reduced matrix expression under loading^87^. Fernando et al. reported loading-frequency-dependent changes in NP-cell ATP content, while Wang et al. showed that SIRT1-dependent mitophagy protected compressed NP cells against mitochondrial injury^55,88^. In this context, the RNM’s concurrent ECM, metabolic, antioxidant, and survival activities describe a mechanically supported environment for NP matrix maintenance.

HL shifted this organization toward matrix remodeling and inflammatory regulation. The RNM combined higher COL1A1 (0.999) with near-maximal MMP13 and ADAMTS4/5 activation and strong NF-*κ*B (0.979 ± 0.033) and AP-1 activity. Zhang et al. found that human NP cells exposed to prolonged compression with shorter rest periods developed greater matrix degradation and NF-*κ*B activation than cells given shorter loading and longer recovery intervals^27^. Wu et al. showed that mechanical stress induced NF-*κ*B-dependent periostin expression, which further reinforced NF-*κ*B signaling and senescence-associated catabolic responses^48^. Sun et al. identified PIEZO1-dependent Ca^2+^/NF-*κ*B signaling upstream of NLRP3 inflammasome activation in NP cells^39^. These findings link the RNM’s HL-associated matrix remodeling to a sustained inflammatory and mechanosensitive regulatory program.

The HL phenotype also involves mitochondrial integrity and survival. ROS activation reached 0.790 ± 0.409, while BAX and CASP3 were elevated and HSP70 was reduced to 0.198 ± 0.378. Hu et al. demonstrated that HSP70 limits compression- induced NP-cell apoptosis by promoting SIRT3 expression, reducing mitochondrial fission and ROS generation^89^. The decline in protective activity within the RNM therefore occurs alongside mechanisms capable of reinforcing oxidative injury and cell death^90^. PI3K-E activation (1.000), contrasting with the nearly inactive PI3K-M branch, places PI3K–AKT-related signaling in a different regulatory setting under HL. TGF*β* (0.986) and YAP/TAZ (0.930) also remained active. Bian et al. found that deficient and excessive integrin-mediated TGF*β* signaling can both impair disc homeostasis, while Wang et al. linked higher hydrostatic pressure to nuclear YAP and NP-cell apoptosis^26,91^. Their context-dependent activity fits the model’s mixed remodeling and injury-associated HL phenotype.

Hypo is particularly informative because reduced mechanical stimulation does not eliminate cellular stress. The RNM retained sustained-calcium activity (Ca^2+^, 0.481 ± 0.455) alongside ROS (0.521 ± 0.493) and BAX (0.530 ± 0.501), even as the mechanically driven calcium branch was largely inactive. Yasuoka et al. found reduced NP glycosaminoglycan content after rat-tail suspension, and Xie et al. showed that spaceflight microgravity impaired PIEZO1-dependent NP mechanosensing and promoted AMPK-associated apoptosis^31,92^. The model’s Hypo AMPK activity (0.594 ± 0.381) is consistent with this form of under-stimulation-associated stress signaling. A possible link to ROS involves diminished cell–matrix survival support: Wu et al. identified fibronectin–integrin/FAK regulation of anoikis-related responses in NP cells, while Kamarajugadda et al. directly measured increased mitochondrial superoxide following matrix detachment in mammary epithelial cells^93,94^. Booth et al. demonstrated that mitochondrial oxidative bursts sensitize ER IP_3_ receptors and promote local Ca^2+^ release; Lin et al. identified ROS-dependent IP_3_R-mediated ER–mitochondrial calcium transfer in compressed NP cells^95,96^. These findings suggest a route by which weaker adhesion-associated signaling could coexist with intracellular calcium release and mitochondrial stress under Hypo. The complete sequence under NP unloading remains a mechanistic hypothesis. The substantial Hypo SOD2/CAT activation (each 0.750 ± 0.390), along with AMPK/ULK1 signaling, suggests that antioxidant and energy-sensing programs remain engaged alongside ROS and sustained calcium, producing a mixed stress-adaptive phenotype.

The close coordination of several outputs also reflects the RNM’s regulatory resolution. Shared upstream inputs can produce near-identical inflammatory and protease activation patterns, while near-zero NL MMP activation represents strong modeled suppression within the specified network; physiological NP matrix turnover still involves basal metalloproteinase activity^97^. Viewed in this context, the three baseline states describe distinct balances between mechanically supported homeostasis, stress adaptation, and injury-associated regulation. Their main contribution is to identify how these processes coexist within each loading regime, providing a network-level interpretation of the simulated NP phenotypes.

### 3.3 Model falsification against experimentally established NP responses

The independent falsification test showed concordance between RNM predictions and 37 of 39 literature-derived NP mechanobi- ological directionality benchmarks (94.9%), including all 6 positive (+) and 31 of 33 negative (−) directionality benchmarks. The agreement was particularly evident in the balance between matrix maintenance and remodeling. Zhang et al. reported that shorter daily periods of compression at 0.8 MPa followed by longer periods at 0.1 MPa maintained a more favorable matrix profile in cultured human NP cells than prolonged exposure to 0.8 MPa, which increased MMP3, MMP13, and ADAMTS4/5 and reduced matrix-associated gene expression^27^. Chen et al. independently observed significantly reduced MMP13 protein expression under moderate fluid shear stress (12 dyn/cm^2^) in rat NP cells^98^. In human non-degenerate NP cells, physiological dynamic hydrostatic pressure increased aggrecan expression significantly, with increase in SOX9 and collagen-II expression^99^. Neidlinger-Wilke et al. observed reduced aggrecan expression in human NP cells at 2.5 versus 0.25 MPa hydrostatic pressure, with reduced collagen-II and increased MMP3 expression also observed in bovine NP cells^32^. Li et al. reported increased TIMP3 expression under lower-frequency compression (0.1–3 Hz), and significant suppression at 5 Hz in immature porcine NP tissue, whereas Huang et al. observed compression-induced ADAMTS4/5 upregulation in human NP cells without a significant change in TIMP3^86,100^. Low hydrostatic pressure reduced collagen-I protein abundance in rat NP cells, whereas sustained compression increased COL1A1 mRNA expression in human NP cells^101,102^. At the tissue level, prolonged upright posture in rats was associated with increased whole-disc COL10A1 and MMP13 expression, and compression of young-adult mouse-tail discs increased RUNX2 transcription^103,104^. The RNM thus reproduced directional changes spanning matrix constituents, protease activity regulators, and markers associated with degenerative tissue remodeling.

Inflammatory responses were also concordant across markers associated with cytokine signaling, chemokine release, and prostaglandin synthesis. Wang et al. showed that 9% and 19% cyclic tensile strain, in contrast to 3% strain, increased TNF-*α*, IL1*β* , and IL6 responses in human NP cells, with high strain also activating NF-*κ*B p65 signaling^105^. Gawri et al. reported increased IL8, CCL2/MCP-1, CXCL1/GRO-*α*, and TGF-*β* 1 secretion from mechanically stretched primary human disc-cell cultures, supporting the RNM’s loading-associated inflammatory and remodeling directions^29^. In rabbit NP cells, Sowa et al. demonstrated that COX-2 transcription depended on compression magnitude and duration^106^. Cambria et al. detected a modest increase in PGE_2_ release 48 h after hyperphysiological compression of bovine NP constructs and investigated its modulation through TRPV4^35^. Easson et al. subsequently showed that static and dynamic disc compression generated different cytokine networks, including distinct TRPV4-dependent relationships among inflammatory and vascular mediators^49^. Agreement across transcriptional responses and secreted mediators is consistent with coordinated inflammatory regulation in the RNM, while the distinct loading-dependent cytokine profiles provide a biological basis for differences between experimental protocols.

The survival and stress-response benchmarks extend this concordance to apoptosis, mitochondrial regulation, and intracellu- lar kinase signaling. Wang et al. demonstrated that low-magnitude cyclic compression (2%) increased Bcl2 and decreased cleaved caspase-3 expression in rat NP cells, whereas high-magnitude compression (20%) produced opposite responses^107^. Zhang et al. also reported increased BAX expression under prolonged compression^27^. High-magnitude compression indepen- dently increased cleaved caspase-3 in 3D rat NP-cell cultures^108^, while sustained 3 MPa compression increased caspase-9 activity in human NP cells^109^. Hu et al. associated compression of human NP-derived stem/progenitor cells with reduced SOD2 expression and increased oxidative injury^110^. In rat NP cells, high-magnitude cyclic compression increased ROS production and p38 phosphorylation relative to low-magnitude compression, while compression-induced ROS was also implicated in JNK-dependent autophagic signaling^84,111^. Kang et al. demonstrated increased DRP1 recruitment to mitochondria during sus- tained compression, linking mitochondrial fission with oxidative stress and apoptosis^112^. Separately, Li et al. observed reduced inhibitory GSK3*β* phosphorylation under high- versus low-magnitude compression and showed that N-cadherin–PI3K/Akt signaling mitigated the associated apoptotic response^113^. These concordant directions cover both early stress signaling and downstream processes governing mitochondrial integrity and NP-cell survival.

The mechanotransductive signaling benchmarks offer a mechanistic connection between the predicted loading inputs and these cellular responses. Shi et al. found that high-magnitude compression increased PIEZO1 expression in human NP cells and altered autophagy–apoptosis regulation^43^. Under 20% cyclic tensile stretch, primary rat NP cells displayed increased PIEZO1 expression, nuclear YAP localization, and TEAD signaling, accompanied by increased COL1 and reduced COL2 expression^114^. In a separate hydrostatic-pressure model, Shi et al. reported that 30 atm increased Wnt3a and *β* -catenin protein expression, with associated matrix-degrading responses^115^. Ke et al. linked sustained compression to actomyosin remodeling, RhoA/ROCK signaling, and MRTF-A nuclear translocation in human NP cells, while Song et al. connected cyclic tensile strain to RhoA–MRTF-A-dependent fibrotic regulation^102,116^. Another compressive-loading study identified PKN1 phosphorylation and downstream KRT8 phosphorylation as regulators of autophagosome initiation^117^. More recently, Mao et al. reported increased angiogenesis-associated expression, including VEGFA, after sustained compression of rat NP cells and identified YAP-condensate disruption as a mechanism contributing to disc vascularization^118^. This latter mechanism differs from the nuclear-YAP response observed during cyclic stretching, indicating that YAP-associated regulation depends on the mechanical and cellular context. The model’s agreement across mechanosensitive channels, transcriptional regulators, and kinase responses is consistent with multiple routes connecting mechanical stimulation to matrix remodeling, autophagy, and vascular-associated signaling.

HSF1 and CCN2 were the two contradictory benchmarks. The RNM predicted higher HSF1 activation under NL, whereas the assigned literature-derived direction was based on HSF1 induction under compression. Chooi and Chan found that HSF1 transcription in bovine NP-cell collagen constructs increased under 30% static strain but decreased under 70% static strain, with responses also varying according to loading and post-loading duration^119^. The disagreement therefore concerns a regulator whose experimental direction is sensitive to strain magnitude and timing, making its loading-dependent control a specific target for further investigation. The CCN2 prediction presents a different regulatory discrepancy: the RNM correctly predicted higher MRTF-A activation under HL but higher CCN2 activation under NL. Song et al. demonstrated that 15% cyclic tensile strain increased nuclear MRTF-A and CCN2/CTGF in human NP cells and that interference with RhoA–MRTF-A signaling attenuated fibrotic responses^116^. Other studies identify additional inputs to CCN2 regulation: SOX9 directly promotes CCN2 transcription in NP cells, while inflammatory cytokines suppress CCN2 and exogenous CCN2 can attenuate IL-1*β* -induced catabolic responses^120,121^. The RNM therefore reproduces the loading direction of MRTF-A but not its experimentally associated CCN2 response, suggesting that the balance among cytoskeletal, matrix-associated, and inflammatory inputs to CCN2 requires closer examination.

The experimental literature also shows why the assignment of a directional loading benchmark requires a well-defined comparator. Studies such as those of Zhang et al. and Wang et al. directly compared graded or differently timed loading regimes, while other studies evaluated excessive loading against unstretched or uncompressed controls^27,105^. Le Maitre et al. examined physiological hydrostatic stimulation in non-degenerate and degenerate NP cells, whereas Neidlinger-Wilke et al. compared two hydrostatic pressures within a physiological range; neither design is mechanically equivalent to an excessive-versus-physiological cyclic-strain experiment^32,99^. Loading magnitude alone is insufficient to define the biological stimulus: the frequency- and duration-dependent changes described by Li et al. and the distinct static and dynamic cytokine responses reported by Easson et al. show that loading history also matters^49,86^. Experimental systems further differ in species, cell maturity, cell culture versus intact-disc organization, and the use of mRNA, total protein, phosphorylation, nuclear localization, or secreted factors as molecular readouts. These differences constrain the availability of directly comparable NP mechanobiology benchmarks and make the experimental loading comparison and measured molecular response essential to the interpretation of each assigned direction.

### 3.4 Mechanistic interpretation of the Hyper-to-Normal rescue screen

The Hyper-to-Normal rescue test examined whether targeted regulatory perturbations could redirect the HL-associated network state toward NL while excessive ML was maintained. The strong ECM response to SOX9 activation (82.8% rescue), with only a modest increase following additional ROS suppression (85.9%), suggests that SOX9-associated regulation accounts for much of the recoverable ECM profile under HL (Fig. 6A). SOX9 also exhibited the second-highest out-degree and ranked among the leading nodes in betweenness centrality, indicating extensive downstream connectivity and a prominent bridging position within the network (Fig. 2). Its predicted influence is supported by experimental studies. Paul et al.^122^ reported increased type II collagen expression following SOX9 delivery to human degenerated disc cells and preservation of NP architecture in rabbit discs. Zhang et al.^123^ observed increased proteoglycan and collagen accumulation in SOX9-transduced bovine NP cells. Tsingas et al.^79^ demonstrated matrix deterioration and disc-cell loss following conditional Sox9 deletion in adult mice. In the RNM, SOX9–ROS increased mean ACAN and COL2A1 activities from 0.216 and 0.252 under HL to 0.986 and 0.994, respectively, while TIMP3 increased from less than 0.001 to 0.716 (Fig. 6A). The TIMP3 response is consistent with the anti-catabolic effects reported following TIMP3 overexpression in NP models^124^, although direct SOX9-dependent regulation of TIMP3 was not established experimentally. The limited additional ECM benefit of other SOX9-containing combinations is illustrated by mTORC1, which ranked highest in betweenness centrality but increased ECM rescue to only 83.8% when suppressed alongside SOX9 activation. This difference indicates that topological prominence does not necessarily translate into greater intervention effectiveness within a given module. The response also differed across functional outputs: SOX9 activation achieved only 15.4% transcription-factor rescue, increasing to 30.7% with ROS suppression (Fig. 6C). The larger incremental effect of ROS suppression in this module (15.3 versus 3.1 percentage points in ECM) suggests that matrix-associated recovery and broader transcriptional regulation differ in their sensitivity to the tested redox perturbation.

The contribution of redox regulation was also apparent in the oxidative-stress defense and proteostasis module, where FOXO activation combined with ROS suppression achieved 68.9% rescue, followed by SIRT1–ROS (60.8%) and NRF2 activation alone (approximately 60.4%) (Fig. 7C). ROS ranked second in betweenness centrality and first in harmonic closeness, reflecting its bridging position and extensive directed reachability within the network (Fig. 2). The substantial recovery associated with different protective regulators suggests that several regulatory mechanisms can influence the oxidative-stress-associated profile under HL^125^. Experimental evidence supports their distinct biological functions. Li et al.^84^ found that high-magnitude cyclic compression (20% versus 2% deformation, 1 Hz, 6 h/day) increased ROS accumulation and senescence in rat NP cells, with both responses attenuated by the ROS scavenger N-acetylcysteine. Alvarez-Garcia et al.^82^ demonstrated impaired antioxidant and autophagy-related regulation following conditional FOXO-family deletion in mouse discs, while Wang et al.^126^ reported reduced mitochondrial dysfunction and apoptosis following FOXO3 overexpression in nutrient-deprived NP cells. Under high-magnitude compression, SIRT1 overexpression protected human NP cells against mitochondrial injury and senescence through PINK1-dependent mitophagy^88^. Tian et al.^127^ similarly demonstrated that bardoxolone methyl, a pharmacological activator of NRF2 signalling, attenuated oxidative injury and apoptosis in compressed human NP cells, with diminished protection following NRF2 silencing. These findings provide a mechanistic basis for the predicted oxidative-stress rescue through FOXO-associated autophagy, SIRT1-dependent mitochondrial quality control, and NRF2-mediated antioxidant defence^128^. Their potentially overlapping protective effects offer a specific basis for investigating whether the RNM-predicted differences among FOXO–ROS, SIRT1–ROS, and NRF2 interventions are reproduced under comparable excessive-loading conditions.

The substantial recovery predicted for oxidative-stress defense did not extend to the same degree to cell-survival-associated regulation. NRF2 activation combined with PIEZO1 suppression achieved the highest rescue in the cell-survival, apoptosis, mitophagy, and DNA-damage module (37.5%), followed by PPAR*γ*–ROS (34.9%) and ROS suppression alone (34.2%) (Fig. 7D). These responses suggest that restoring antioxidant and proteostasis-associated regulation may be insufficient to normalize the broader cell-survival profile under HL. The involvement of PIEZO1 is consistent with experimental evidence linking mechanosensitive signalling to compression-induced cellular injury^129,130^. Shi et al.^43^ found that high-magnitude cyclic compression (15% versus 1.5% deformation) increased PIEZO1 expression, impaired autophagy, and promoted apoptosis and matrix degradation in human NP cells. Treatment with GsMTx4, a mechanosensitive ion-channel inhibitor that also inhibits PIEZO1, partially attenuated these effects. Li et al.^131^ further reported reduced apoptosis and matrix injury following PIEZO1 deletion or pharmacological inhibition in mouse disc-degeneration models. In human NP-derived mesenchymal stem cells subjected to 1.0 MPa compression, the PPAR*γ* agonist pioglitazone reduced ROS accumulation, mitochondrial injury, and apoptosis^80^. These findings provide a biological basis for evaluating the predicted NRF2–PIEZO1 and PPAR*γ*–ROS combinations, particularly whether coordinated modulation of mechanosensitive injury and cytoprotective signalling improves apoptosis and mitochondrial homeostasis under excessive loading.

Inflammatory and proteolytic regulation showed a greater response to combined perturbations than to individual target modulation. SOX9 activation and NF-*κ*B suppression achieved 17.9% and 3.6% rescue, respectively, whereas their combination increased rescue to 57.0% (Fig. 6D). Similarly, combining PPAR*γ* activation with NF-*κ*B suppression increased rescue from 17.6% to 46.2%. The SOX9–NF-*κ*B intervention reduced mean IL8, CCL2, CXCL1, and CXCL3 activities from approximately 1.000 under HL to 0.008, and MMP1, MMP3, and MMP13 from approximately 0.999 to below 0.001 (Fig. 6A). NF-*κ*B exhibited the highest out-degree (24 outgoing interactions) and second-highest harmonic closeness in the network (Fig. 2), consistent with its extensive connections to inflammatory and catabolic regulation. Its limited single-node rescue, however, suggests that NF-*κ*B suppression alone does not adequately redirect the broader module toward NL, whereas concurrent SOX9 or PPAR*γ* activation produces a substantially greater response. Experimental evidence supports the involvement of NF-*κ*B in mechanically induced matrix catabolism^132^. Glaeser et al.^133^ investigated NF-*κ*B pathway inhibition using a NEMO- binding-domain (NBD) peptide in IL-1*β* -stimulated human NP cells cultured in fibrin constructs under cyclic compression (5 kPa, 1 Hz, 48 h). NBD treatment reduced MMP3 expression and release and improved cell viability, although IL8 was not significantly reduced. In a rabbit disc-puncture model, Zhao et al.^134^ demonstrated greater preservation of collagen-II and proteoglycan-associated outcomes following combined SOX9 overexpression and MMP3 knockdown than with either intervention alone. These studies provide experimental support for anti-catabolic regulation through NF-*κ*B inhibition and complementary anabolic reinforcement. The broader chemokine and protease suppression predicted for both SOX9–NF-*κ*B and PPAR*γ*–NF-*κ*B identifies an experimentally testable extension of these findings under excessive mechanical loading.

The SOX9–NF-*κ*B combination also achieved 66.0% rescue in the growth-factor module, compared with 20.7% for SOX9 activation alone (Fig. 6B). This response suggests that inflammatory regulation influences the HL-associated growth- factor profile in the RNM. Experimental evidence supports interactions between these regulatory processes^135^. Shen et al.^136^ demonstrated that GDF5 overexpression in lipopolysaccharide-treated rat NP cells reduced inflammatory mediator production and NF-*κ*B activation. In mouse lumbar-instability and notochordal-cell-specific knockout models, Bian et al.^91^ found that both excessive integrin-mediated TGF-*β* activation and disruption of TGF-*β* signalling compromised disc homeostasis. These studies demonstrate the context-dependent regulation of growth-factor pathways, although they do not directly establish the predicted SOX9–NF-*κ*B response. The phenotype-panel results further distinguish the effects of the leading combinations. Despite similar ECM rescue with SOX9–ROS and SOX9–NF-*κ*B (85.9% and 85.2%, respectively), SOX9–NF-*κ*B produced higher mean anabolic activity (0.718 versus 0.589) and lower catabolic activity (0.371 versus 0.653) (Supplementary Table S4). These differences indicate that comparable ECM-profile recovery can accompany distinct responses across the wider anabolic and catabolic network. Experimental assessment of these combinations would therefore benefit from including growth-factor, inflammatory, and proteolytic readouts alongside conventional ECM markers.

The present study is subject to several limitations related to network construction, regulatory representation, and simulation assumptions. The curated RNM comprises 146 nodes and 353 literature-supported interactions, representing selected compo- nents of NP-cell mechanotransduction without encompassing all molecules and regulatory interactions involved in this complex biological process. Although NP-specific experimental evidence was prioritized, some interactions were informed by studies in related mechanosensitive cells. Regulatory interactions were represented as activating or inhibitory relationships without explicitly describing the underlying molecular processes, such as phosphorylation, protein complex formation, or nuclear translocation. The SQUADS formalism employs normalized, non-parametrically specified regulatory constants, capturing network-level behaviour without experimentally calibrated kinetic parameters. Mechanical stimulation was represented by discrete Hypo, NL, and HL inputs, and network responses were evaluated at steady state under fixed loading conditions, limiting the assessment of transient signalling, loading history, and time-dependent adaptation. The rescue test examined 48 predefined single- and dual-node interventions, modeled as sustained activation or suppression of selected nodes without considering intermediate activity levels, other target combinations, or time-dependent intervention strategies. Rescue scores quantify the percentage reduction in Euclidean distance between ensemble-mean module profiles and the NL reference, relative to the baseline HL-to-NL distance, and therefore depend on module composition and baseline separation. These scores describe recovery of regulatory profiles within the RNM and cannot be equated with corresponding percentages of cellular or tissue recovery.

Overall, the RNM provides a mechanistic framework for interpreting NP-cell responses to mechanical loading and prioritizing experimentally testable regulatory interventions. Future work should focus on incorporating additional NP-specific regulatory evidence, graded and time-dependent loading, and experimentally calibrated parameters to improve the model’s mechanistic and quantitative reliability. Experimental evaluation of the predicted SOX9–ROS, SOX9–NF-*κ*B, FOXO–ROS, and NRF2–PIEZO1 interventions, together with targeted investigation of the contradictory HSF1 and CCN2 responses, will be important for assessing the biological relevance of the model predictions and their potential translational applicability.

## 4 Materials and Methods

### 4.1 Regulatory network construction and structural analysis

Regulatory network modeling (RNM) was employed to represent NP-cell mechanotransduction as a qualitative, signed directed graph, in which nodes correspond to molecular regulators or cellular processes and edges encode activating or inhibitory interactions^137,138^. The model integrates primary mechanosensors, intracellular signaling cascades, transcriptional regulators, redox mediators, growth-factor pathways, and extracellular matrix effectors to capture multilevel load-dependent signal integration.

Mechanotransductive interactions were systematically curated from more than 130 peer-reviewed publications using structured searches across PubMed, Scopus, Web of Science, and Google Scholar. Searches combined tissue-specific terms with mechanobiological terms using Boolean operators, including combinations such as ("nucleus pulposus" OR "intervertebral disc") AND ("mechanotransduction" OR "mechanical loading" OR "mechanobiology"). For targeted pathway curation, searches were further refined by adding the name of the molecular node of interest. When NP-specific evidence was limited, relevant findings from related mechanosensitive cells such as chondrocytes were considered if the regulatory direction was conserved and the interacting nodes were independently supported as expressed and functional in NP cells. Curated pathway and interaction databases (Reactome, KEGG, and STRING) were additionally consulted to cross-validate regulatory relationships and identify missing intermediates. Inclusion required direct experimental evidence of mechanical modulation. The complete search strings and knowledge-driven network construction workflow are provided in Supplementary Material.

All regulatory interactions were encoded as activating or inhibitory according to experimental evidence. The resulting network comprises 146 nodes and 353 curated interactions, each annotated with its source and target molecules, regulatory sign and supporting references and, where reported, the cell type, experimental model and culture conditions. These data were compiled within a single structured corpus provided as Supplementary Data 1. The network definition used in all simulations is archived on Zenodo^139^. For topological analysis, the network was represented as a signed directed graph *G* = (*V, E*), where *V* denotes the set of molecular nodes and *E* the directed regulatory interactions between them. The regulatory sign of each interaction was represented by the signed adjacency matrix *A_s_* = [*A_i j_*] (Eq. (1)),

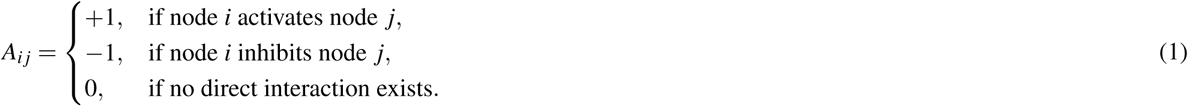

Network visualization and global topological analysis were performed using Cytoscape^73^. The global properties examined included average node degree, network diameter, characteristic path length, network density, clustering coefficient, heterogeneity, and centralization. To characterize the topological position of individual nodes within the directed network, we calculated betweenness centrality (Eq. (2)) and harmonic closeness centrality (Eq. (3)). Betweenness centrality quantifies the extent to which a node lies on shortest directed paths connecting other node pairs and was calculated as

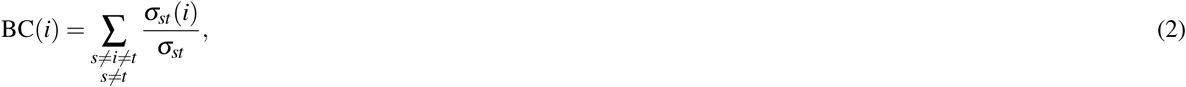

where *σ_st_* is the number of shortest directed paths from node *s* to node *t*, and *σ_st_*(*i*) is the number of those paths that pass through node *i*. Nodes with high betweenness therefore occupy positions that connect otherwise separated parts of the regulatory network and may act as topological intermediates between signaling modules. Harmonic closeness centrality was used to quantify the directed reachability of each node,

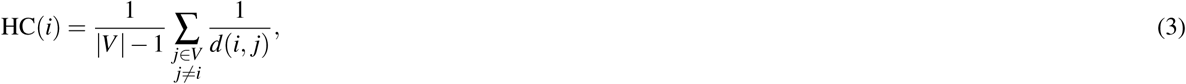

where *d*(*i, j*) is the length of the shortest directed path from node *i* to node *j*, with unreachable nodes contributing zero to the sum. Higher HC(*i*) therefore indicates that a node can reach a larger fraction of the network through relatively short directed paths. To facilitate systematic analysis and interpretation of network-level responses, nodes were further grouped according to established biological functions into ECM-related markers, growth factors, transcription factors, inflammatory mediators, metabolic regulators, ion-channel signaling components, oxidative stress responses, and cell-fate modules.

### 4.2 Encoding mechanical loading (ML) inputs and dynamical formalism

ML was encoded using three external input nodes representing hypo-loading (Hypo), normal loading (NL), and hyper/excessive loading (HL). Each input node was defined over a normalized domain [0, 1]. These inputs do not correspond to direct normalizations of measured physical quantities, such as stress magnitude, loading frequency, or osmolarity, but rather represent qualitative loading regimes derived from experimentally interpreted conditions. Interactions were assigned according to the loading context in which they were experimentally observed, with interactions reported under physiological loading associated with NL and those observed under excessive/sustained or injurious loading associated with HL. Representative experimental loading conditions and their corresponding interpretation as NL or HL are summarized in Table 1. Hypo was introduced as an explicit input to represent reduced mechanical stimulation relative to physiological loading.

Mechanistically, Hypo reduces activation of key mechanosensors, namely integrins and mechanosensitive ion channels, thereby weakening integrin-mediated adhesion signaling such as FAK–PI3K–AKT signaling, and reducing cytoskeletal tension. The resulting loss of mechano-adhesive coupling compromises cell–ECM engagement and pro-survival signaling, promoting apoptosis in a manner consistent with an anoikis-like response^144–147^. NL represents physiological mechanical stimulation that supports matrix homeostasis and anabolic cellular responses. In intervertebral disc mechanobiology, physiological dynamic loading is commonly associated with compressive or hydrostatic magnitudes of approximately 0.1–0.6 MPa applied at 0.1–1 Hz^22,76^. Under this regime, NL predominantly activates *α*_5_*β*_1_ integrin in its intact fibronectin-binding configuration (*α*_5_*β*_1_–FN)^23^, *α_V_ β*_6_ integrin^91^, and the osmo-mechanosensitive channel TRPV4^101^. NP cells also reside within a mechanically coupled hyperosmotic niche of approximately 400–600 mOsm/kg H_2_O under physiological conditions^148^, such that osmotic cues contribute to mechanotransduction under this loading regime^149^. HL represents excessive or injurious mechanical stimulation. Such conditions are commonly associated with higher loading magnitudes or with sustained or otherwise damaging loading patterns that promote inflammatory and catabolic responses^22,86,150^. Under this regime, HL primarily engages *α*_5_*β*_1_ integrin in its fragmented-fibronectin-binding configuration (*α*_5_*β*_1_–Fs)^151^, *α_V_ β*_3_ integrin^121^, and the strain-sensitive channel PIEZO1^39,48^. These assignments represent dominant but not exclusive sensor–load relationships, as mechanosensor activation remains context-dependent and is influenced by matrix composition, osmotic conditions, and cellular phenotype^35,49,152^. The principal mechanosensors and their downstream signaling organization, including cytoskeletal, kinase, redox, inflammatory, transcriptional, and extracellular matrix regulatory modules, are summarized schematically in Figure 8.

**Figure 8.**
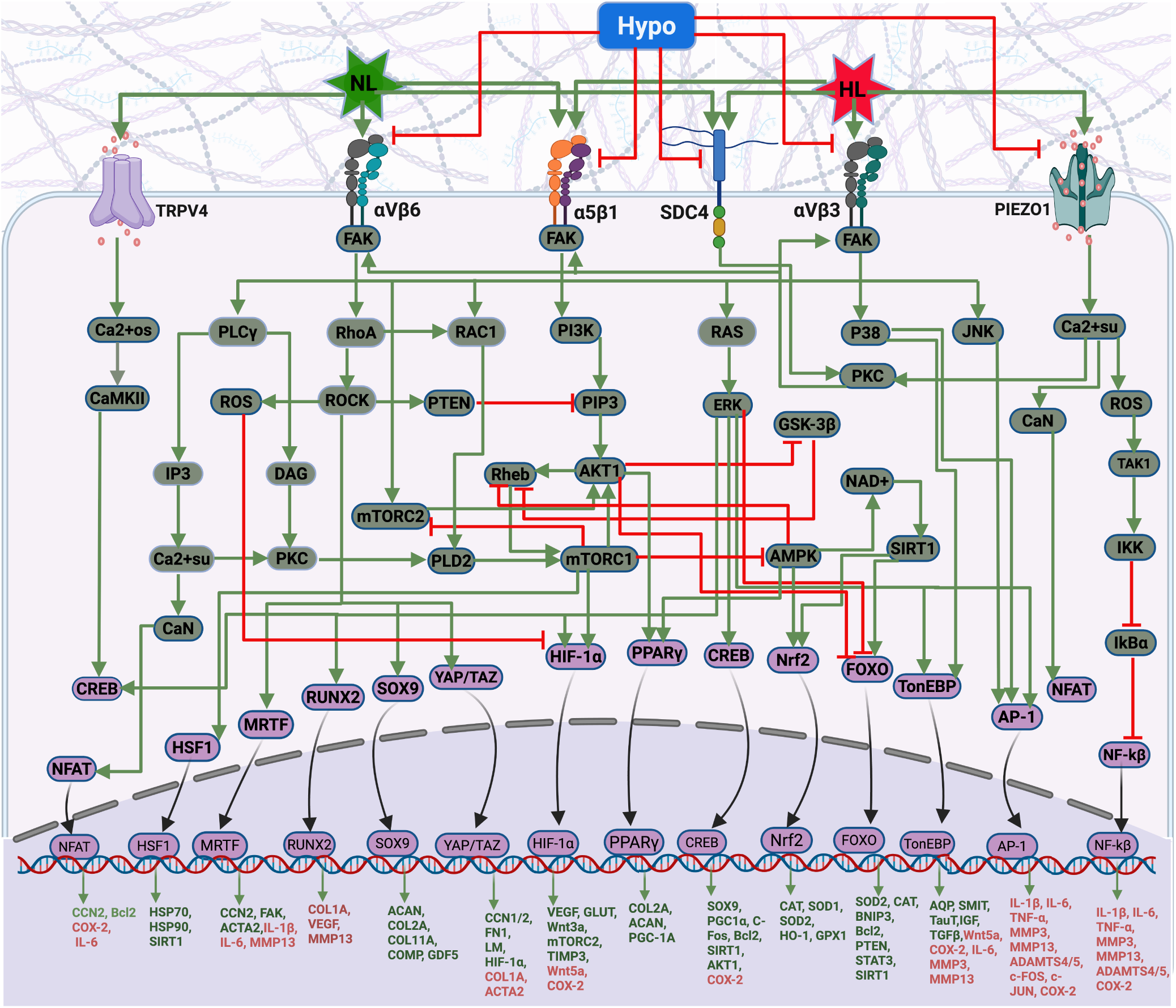
Integrated mechanotransductive signaling pathways in NP cells. The schematic summarizes primary mechanosensors (*α*_5_*β*_1_, *α_V_ β*_3_, *α_V_ β*_6_, SDC4, PIEZO1, and TRPV4) and representative downstream signaling cascades linking mechanical cues to transcriptional regulation and extracellular matrix responses. Green arrows and red T-bars represent activating and inhibitory interactions, respectively. Gene labels in green denote predominantly anabolic transcriptional targets, whereas red labels indicate pro-catabolic or inflammatory targets encoded within the regulatory network model.

To simulate network dynamics, we employed the *Standardized Qualitative Dynamical Systems* (SQUADS) formalism^153,154^, which provides a continuous ODE-based extension of logical regulatory models. Each node *x _j_*(*t*) ∈ [0, 1] represents the normalized activation level of molecule *j* and evolves according to:

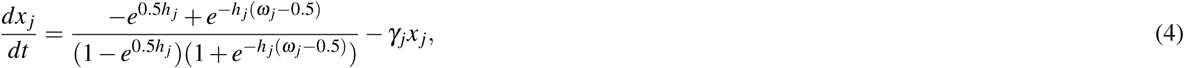

where *h _j_ >* 0 controls the steepness of the activation function (slope at inflexion point), *γ _j_ >* 0 represents passive decay toward baseline, and *ω_j_*, the stimulation, is the effective regulatory input derived from activating and inhibitory signals. The aggregated activating (*A_j_*) and inhibitory (*I_j_*) contributions are defined as

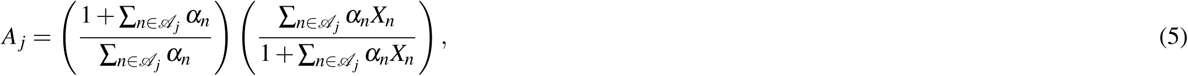

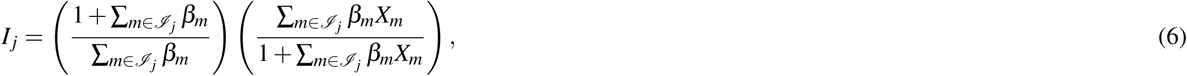

and the effective regulatory input is given by

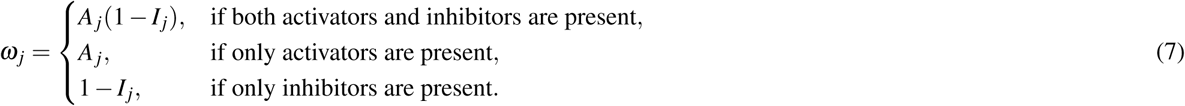

Unless otherwise specified, we set *α_n_*= *β_m_*= *γ _j_* = 1 and *h _j_* = 10^153^. The system was implemented in `MATLAB R2024b` and solved using `ode45` with tolerances `RelTol`= 10^−8^, `AbsTol`= 10^−10^, and `MaxStep`= 0.5. The original MATLAB scripts, an open-source Python reimplementation and an SBML export of the model are publicly available^139,155^ (see Code availability). Simulations were performed over *t* ∈ [0, 100], and steady states were identified when |*dx_j_/dt*| *<* 10^−8^ for all nodes. In baseline simulations, each loading regime was implemented by selective activation of a single input above baseline, while the remaining inputs were fixed at a low baseline value (0.01). Specifically, normal loading was represented by NL = 0.8 (Hypo = HL = 0.01), hyper-loading by HL = 0.8 (Hypo = NL = 0.01), and hypo-loading by Hypo = 0.2 (NL = HL = 0.01). These values were chosen to ensure strong activation of the dominant loading input while maintaining minimal background activity in the non-dominant inputs.

### 4.3 Model falsification using independent NP mechanobiology experiments

To evaluate the ability of the RNM to reproduce experimentally reported mechanobiological responses of NP cells to ML, model-predicted differences between NL and HL conditions were compared with directional molecular responses documented in published experimental studies. The assessment was falsification-based, examining whether RNM predictions were consistent with, or contradicted, literature-derived experimental benchmarks, following the principles of biological model validation and invalidation^156,157^.

Experimental benchmarks were assembled from peer-reviewed and published studies investigating molecular responses of NP cell or intervertebral disc tissue cultures to ML^106,158^. The benchmark panel was curated according to whether the experimental studies reported increased molecular expression or activity under physiological (NL-like) or excessive (HL-like) ML conditions. The classification of experimental loading conditions as NL-like or HL-like was based on the reported mechanical loading parameters and their interpretation in the context of NP mechanobiology. Individual experimental observations were interpreted relative to their reported comparators, which could be unloaded controls or alternative loading conditions. The resulting panel comprised 39 markers with 6 markers reported to exhibit higher expression or activity under NL-like loading conditions and 33 markers reported to exhibit higher expression or activity under HL-like loading conditions. The complete experimental benchmark panel, including the observed molecular responses, NL-like and HL-like loading classifications, mechanical loading conditions and experimental comparators, literature-derived marker-level directionality, and supporting references, is presented in Table 2. The evaluation procedure is summarized in Fig. 9.

**Figure 9.**
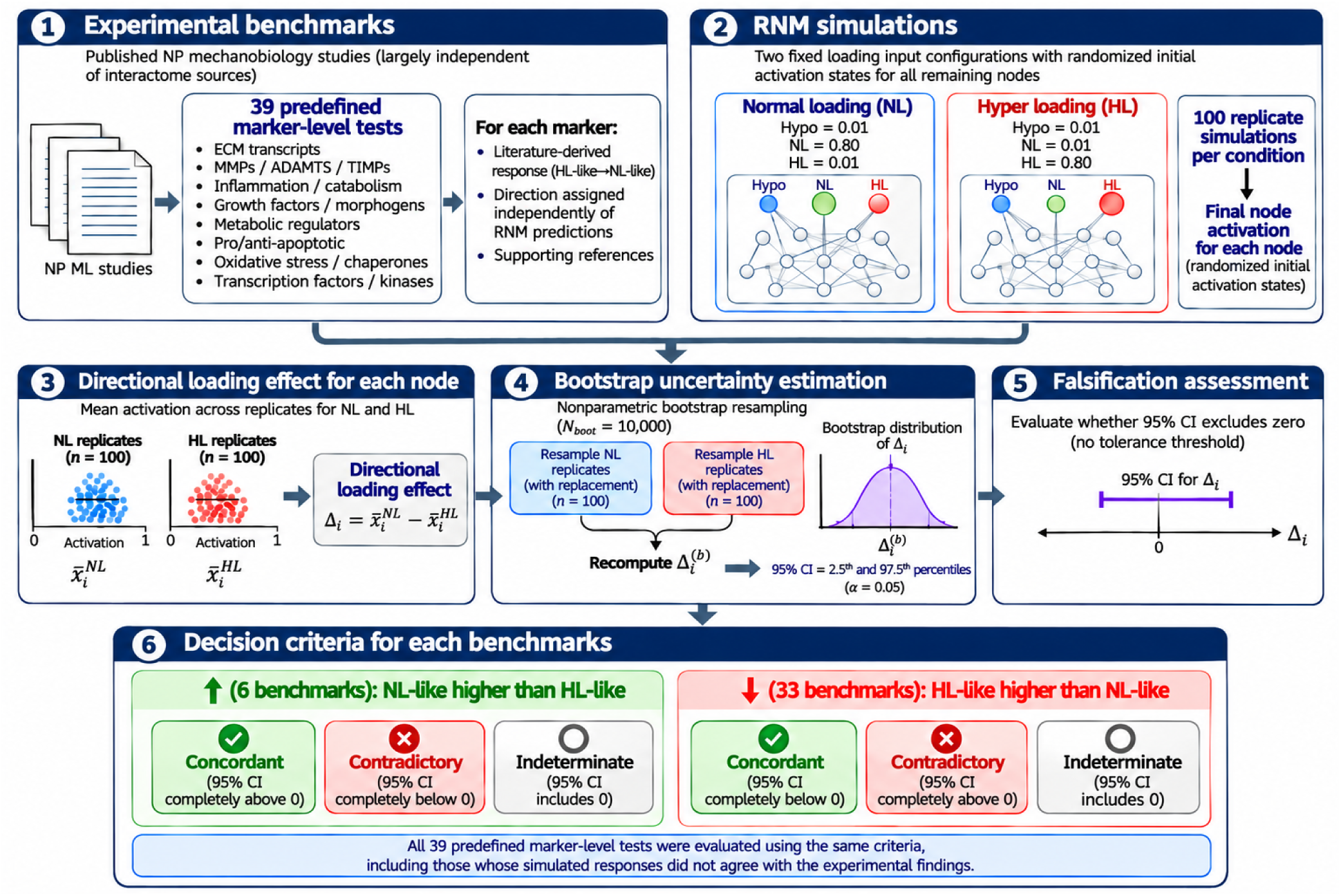
Workflow for evaluating 39 literature-derived directional benchmarks against RNM predictions using 95% bootstrap confidence intervals and directional agreement criteria.

The RNM was simulated under two fixed loading-input configurations: NL (Hypo = 0.01, NL = 0.80, HL = 0.01) and HL (Hypo = 0.01, NL = 0.01, HL = 0.80). For each condition, 100 replicate simulations were performed with randomized initial activation states for all remaining nodes. The final node activations obtained from these simulations were used to quantify the directional loading effect for each benchmark node *i*:

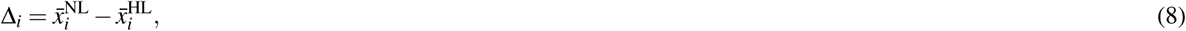

where 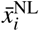 and 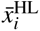 denote the mean final activation of node *i* across replicate simulations under NL and HL, respectively.

Positive values of Δ*_i_*indicate higher mean activation under NL, whereas negative values indicate higher mean activation under HL. Uncertainty in the estimated loading effect was quantified using nonparametric bootstrap resampling^159,160^. For each benchmark node, *N*_boot_ = 10,000 bootstrap iterations were performed by independently sampling with replacement from the NL and HL replicate outcomes, drawing 100 observations from each condition per iteration. The mean activation difference 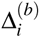 was recalculated for each bootstrap sample. The corresponding 95% bootstrap confidence interval (CI) was defined by the empirical 2.5th and 97.5th percentiles of the bootstrap distribution (*α* = 0.05). These intervals quantify sampling uncertainty in the estimated mean loading effects arising from the finite ensemble of randomized initial conditions. The literature-derived directionality (+ or −) specified in the final column of Table 2 provided the experimental reference against which the direction of each RNM-predicted activation difference (Δ*_i_*) was compared. The 95% bootstrap CI of each Δ*_i_* was used to determine whether the predicted difference was statistically distinguishable from zero and to establish its direction. For the 6 markers with positive literature-derived directionality (+), predictions were classified as *concordant* when the lower CI bound exceeded zero and *contradictory* when the upper CI bound was below zero. Conversely, for the 33 markers with negative literature-derived directionality (−), predictions were classified as *concordant* when the upper CI bound was below zero and *contradictory* when the lower CI bound exceeded zero. Predictions were classified as *indeterminate* when the CI included zero, indicating that the RNM simulations did not establish a definitive direction of the activation difference. All 39 benchmarks were evaluated using the same CI-based directional classification criteria.

### 4.4 Hyper-to-Normal rescue screening via node clamping

The Hyper-to-Normal rescue test was designed to evaluate whether targeted perturbations of regulatory activity could shift an HL-associated network state toward the physiological NL reference while the HL input was maintained. Selection of intervention targets combined two complementary forms of evidence. First, experimentally observed responses to target inhibition, activation, or genetic manipulation were used to define biologically motivated perturbation directions, prioritizing studies of NP cells or disc tissue exposed to excessive mechanical loading and extending to inflammatory, nutrient-deprivation, and aging-related injury models where necessary. The experimental systems, interventions, and principal findings supporting these choices are summarized in Table 3.

**Table 3.**
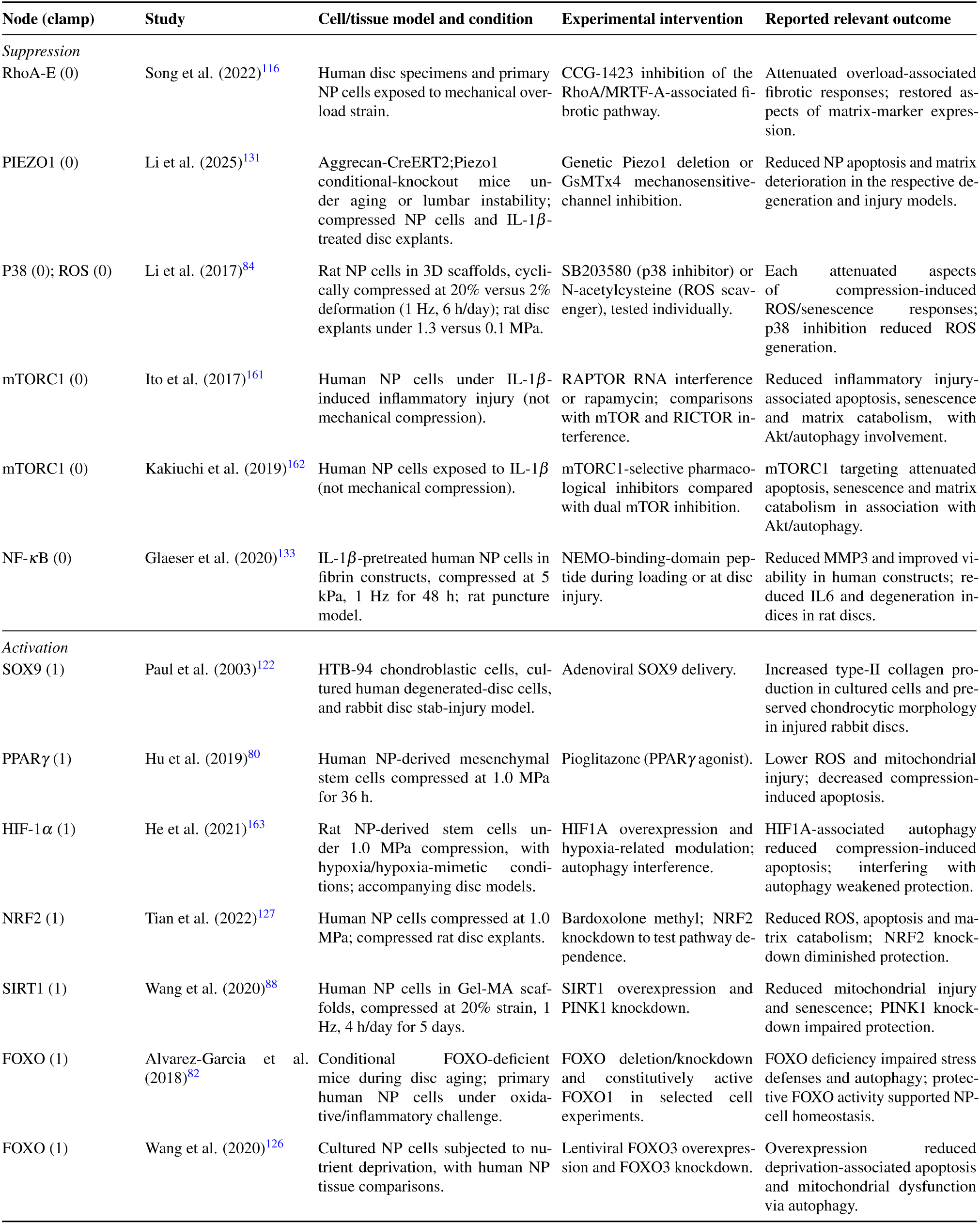
Experimental basis for candidate node-clamping directions. The studies used different models and perturbations; the clamp values are idealized regulatory interventions, not experimentally matched drug concentrations or effects.

| Node (clamp) | Study | Cell/tissue model and condition | Experimental intervention | Reported relevant outcome |
|---|---|---|---|---|
| *Suppression* |  |  |  |  |
| RhoA-E (0) | Song et al. (2022)<sup>116</sup> | Human disc specimens and primary NP cells exposed to mechanical over-load strain. | CCG-1423 inhibition of the RhoA/MRTF-A-associated fi-brotic pathway. | Attenuated overload-associated fibrotic responses; restored as-pects of matrix-marker expres-sion. |
| PIEZO1 (0) | Li et al. (2025)<sup>131</sup> | Aggrecan-CreERT2;Piezo1 conditional-knockout mice un-der aging or lumbar instability; compressed NP cells and IL-1β-treated disc explants. | Genetic Piezo1 deletion or GsMTx4 mechanosensitive-channel inhibition. | Reduced NP apoptosis and matrix deterioration in the respective de-generation and injury models. |
| P38 (0); ROS (0) | Li et al. (2017)<sup>84</sup> | Rat NP cells in 3D scaffolds, cycli-cally compressed at 20% versus 2% deformation (1 Hz, 6 h/day); rat disc explants under 1.3 versus 0.1 MPa. | SB203580 (p38 inhibitor) or N-acetylcysteine (ROS scav-enger), tested individually. | Each attenuated aspects of compression-induced ROS/senescence responses; p38 inhibition reduced ROS generation. |
| mTORC1 (0) | Ito et al. (2017)<sup>161</sup> | Human NP cells under IL-1β-induced inflammatory injury (not mechanical compression). | RAPTOR RNA interference or rapamycin; comparisons with mTOR and RICTOR in-terference. | Reduced inflammatory injury-associated apoptosis, senescence and matrix catabolism, with Akt/autophagy involvement. |
| mTORC1 (0) | Kakiuchi et al. (2019)<sup>162</sup> | Human NP cells exposed to IL-1β (not mechanical compression). | mTORC1-selective pharmaco-logical inhibitors compared with dual mTOR inhibition. | mTORC1 targeting attenuated apoptosis, senescence and matrix catabolism in association with Akt/autophagy. |
| NF-κB (0) | Glaeser et al. (2020)<sup>133</sup> | IL-1β-pretreated human NP cells in fibrin constructs, compressed at 5 kPa, 1 Hz for 48 h; rat puncture model. | NEMO-binding-domain pep-tide during loading or at disc injury. | Reduced MMP3 and improved vi-ability in human constructs; re-duced IL6 and degeneration in-dices in rat discs. |
| *Activation* |  |  |  |  |
| SOX9 (1) | Paul et al. (2003)<sup>122</sup> | HTB-94 chondroblastic cells, cul-tured human degenerated-disc cells, and rabbit disc stab-injury model. | Adenoviral SOX9 delivery. | Increased type-II collagen pro-duction in cultured cells and pre-served chondrocytic morphology in injured rabbit discs. |
| PPARγ (1) | Hu et al. (2019)<sup>80</sup> | Human NP-derived mesenchymal stem cells compressed at 1.0 MPa for 36 h. | Pioglitazone (PPARγ agonist). | Lower ROS and mitochondrial injury; decreased compression-induced apoptosis. |
| HIF-1α (1) | He et al. (2021)<sup>163</sup> | Rat NP-derived stem cells un-der 1.0 MPa compression, with hypoxia/hypoxia-mimetic condi-tions; accompanying disc models. | HIF1A overexpression and hypoxia-related modulation; autophagy interference. | HIF1A-associated autophagy reduced compression-induced apoptosis; interfering with autophagy weakened protection. |
| NRF2 (1) | Tian et al. (2022)<sup>127</sup> | Human NP cells compressed at 1.0 MPa; compressed rat disc explants. | Bardoxolone methyl; NRF2 knockdown to test pathway de-pendence. | Reduced ROS, apoptosis and ma-trix catabolism; NRF2 knock-down diminished protection. |
| SIRT1 (1) | Wang et al. (2020)<sup>88</sup> | Human NP cells in Gel-MA scaf-folds, compressed at 20% strain, 1 Hz, 4 h/day for 5 days. | SIRT1 overexpression and PINK1 knockdown. | Reduced mitochondrial injury and senescence; PINK1 knock-down impaired protection. |
| FOXO (1) | Alvarez-Garcia et al. (2018)<sup>82</sup> | Conditional FOXO-deficient mice during disc aging; primary human NP cells under oxida-tive/inflammatory challenge. | FOXO deletion/knockdown and constitutively active FOXO1 in selected cell experiments. | FOXO deficiency impaired stress defenses and autophagy; protec-tive FOXO activity supported NP-cell homeostasis. |
| FOXO (1) | Wang et al. (2020)<sup>126</sup> | Cultured NP cells subjected to nu-trient deprivation, with human NP tissue comparisons. | Lentiviral FOXO3 overexpres-sion and FOXO3 knockdown. | Overexpression reduced deprivation-associated apoptosis and mitochondrial dysfunction via autophagy. |

Second, the structural analysis described in Section 4.1 was used to assess each candidate’s position in the curated directed network. Signed out-degree was considered as an indicator of direct downstream connectivity, betweenness centrality as an indicator of potential bridging between network regions, and harmonic closeness as a measure of directed reachability. These measures provided complementary topological context for prioritizing regulators with potentially distributed downstream effects; they were not treated as evidence of therapeutic efficacy or as a numerical threshold for inclusion. The screen retained experimentally motivated candidates with distinct mechanistic functions, irrespective of whether they ranked highly for all three topology metrics. Guided by this, six regulatory nodes were selected for suppression (RhoA-E, PIEZO1, ROS, P38, mTORC1, and NF-*κ*B), each clamped to 0, and six for activation (SOX9, PPAR*γ*, HIF-1*α*, NRF2, SIRT1, and FOXO), each clamped to 1. The *in silico* node-clamping screen comprised 12 single-node and 36 dual-node interventions, with each dual intervention combining one suppression target and one activation target, giving 48 interventions in total.

The Hypo, NL, and HL baseline ensembles were generated independently from 100 randomized initial conditions under the corresponding fixed loading inputs. The NL ensemble defined the reference state. Each intervention replicate was initialized from the final steady state of its corresponding HL replicate. At intervention onset, the selected node or nodes were assigned the prescribed values; non-clamped nodes retained their replicate-specific HL activities. The HL loading inputs and intervention clamps remained fixed during the subsequent simulation, while the other nodes evolved according to the governing ODE system.

Rescue was evaluated at three levels: (i) node-resolved changes between paired HL and perturbed steady states; (ii) phenotype-level responses assessed using predefined anabolic and catabolic readout panels; and (iii) recovery of functional- module profiles toward NL, quantified by Euclidean distance between ensemble-mean activation vectors. The anabolic panel comprised ACAN, COL2A1, TIMP3, PPAR*γ*, GDF5, IGF1, SOX9, and Bcl2, representing matrix regulation, growth-factor- associated signalling, and cell survival^80,124,136,164–166^. The catabolic panel comprised MMP1, MMP3, MMP13, ADAMTS4/5, IL1*β* , IL6, IL8, TNF, iNOS, and COX-2, representing matrix proteolysis and inflammatory responses^29,35,97,136,167,168^. Node-resolved and phenotype-level calculations are provided in Supplementary Materials.

For each predefined functional module *C* , the ensemble-mean steady-state activation vectors under HL, NL, and intervention *p* were denoted by 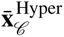, 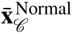, and 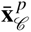, respectively. The baseline HL-to-NL distance and the distance remaining after intervention were defined as

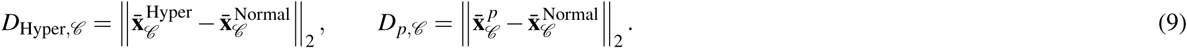

For modules with *D*_Hyper*,C*_ *>* 0, the percentage reduction in profile distance was calculated as

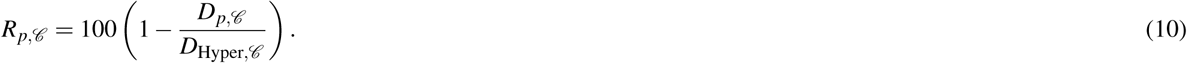

A score of 100% indicates that the perturbed and NL ensemble-mean profiles coincide within the selected module, 0% indicates no reduction in the HL-to-NL distance, and a negative score indicates an increased distance from NL. The 20 highest-scoring interventions were ranked and presented separately for each functional module.

## Author contributions

Z.G.W. conceived the study, curated the regulatory network, developed the model, implemented and performed the simulations and analyses, and drafted the manuscript. F.K.C. developed the open-source Python reimplementation, the SBML export and the web application, and verified the reproducibility of the reported results. J.N. contributed to the conceptualization and overall design of the study, supervised the project, secured funding, provided scientific guidance throughout model development and analysis, contributed to the interpretation of the results, and revised the manuscript. All authors reviewed and approved the final manuscript.

## Data availability

The regulatory network used in all simulations (146 nodes and 353 signed interactions) and the simulation outputs underlying the reported results are archived on Zenodo^139^. The literature-annotated interaction corpus is provided as Supplementary Data 1 (*NP-MT-RNM_Supplementary_Data_1_Interaction_Corpus.xlsx*).

## Code availability

The original MATLAB scripts, an open-source Python reimplementation that regenerates the analyses and figures reported here, and an SBML Level 3 Version 2 export of the model are archived on Zenodo^139^ and maintained at https://github.com/SpineView1/np-mt-rnm. An interactive web application for exploring the model, simulating user-defined loading conditions and node clamps, and browsing the reported results is archived on Zenodo^155^ and maintained at https://github.com/SpineView1/np-mt-rnm-web.

