## Supplementary Materials for "A systems-level view of nucleus pulposus mechanotransduction and its regulatory control"

### ABSTRACT

Intervertebral disc degeneration (IDD), a major contributor to chronic low back pain, is strongly influenced by mechanical loading-dependent regulation of nucleus pulposus (NP) cell phenotype. Previous studies have characterized individual mechanosensors and specific signalling pathways involved in NP mechanotransduction. However, the regulatory interactions through which these components integrate mechanical cues into coordinated cellular responses remain incompletely understood. Here, we present an NP-specific, signed, directed regulatory network model (RNM) comprising 146 molecular nodes and 353 literature-supported interactions. The network connects mechanosensors with signalling cascades, metabolic and redox regulators, transcription factors, extracellular matrix (ECM) effectors, inflammatory mediators, and cell-fate pathways. Using the SQUADS semi-quantitative formalism, which accommodates limited kinetic parameter availability, we simulated steady-state responses to hypo-loading (Hypo), normal loading (NL), and hyper-loading (HL). We also analyzed network topology and screened 48 single- and dual-node perturbations under sustained HL. NL maintained high activity of the ECM-associated regulators ACAN, COL2A1, SOX9, and TIMP3, supported by AKT1, FAK, and antioxidant regulation. HL promoted inflammatory and catabolic regulation through mediators such as NF-$\kappa$B, AP-1, MMP13, and ADAMTS4/5, accompanied by increased ROS, BAX, and CASP3 activity. Hypo reduced anabolic activity while retaining calcium, ROS, AMPK, and apoptotic signalling, suggesting a possible adhesion-dependent stress mechanism. Model falsification against 39 literature-derived molecular directionality benchmarks yielded 37 concordant predictions (94.9%), providing indirect, direction-level support for the RNM. HSF1 and CCN2 showed contradictory responses. Simulated regulatory recovery was module-specific. SOX9 activation achieved 82.8% ECM-profile recovery, increasing to 85.9% with ROS suppression. SOX9 activation combined with NF-$\kappa$B suppression achieved 57.0% inflammatory/proteolytic-module recovery. FOXO activation with ROS suppression achieved 68.9% oxidative-stress/proteostasis-module recovery. NRF2 activation with PIEZO1 suppression achieved 37.5% cell-survival-module recovery. These predictions identify candidate molecular targets and intervention combinations for anabolic reinforcement, inflammatory suppression, redox protection, and mechanosensory modulation. The RNM provides a mechanistic framework for computational prioritization of candidate therapeutic targets and intervention combinations. Its predictions can guide subsequent preclinical validation in NP-cell and disc-explant models under controlled mechanical loading. Experimental validation, incorporation of time-dependent loading, and calibration of regulatory parameters are necessary to establish the therapeutic relevance of these predictions and improve the model's translational applicability.

### Supplementary Material

### 1 Method

### Definitions of nodes included in the NP mechanotransduction regulatory network

**Table S1.** Definition of nodes included in the NP mechanotransduction regulatory network. Each node represents a molecular species, molecular complex, biological process, or external input used in the regulatory network model. Nodes with suffixes –*M* and –*E* represent activity under Normal and Hyper mechanical loading, respectively.

| Node ID | Full name | Node type | Functional category | Notes |
|---|---|---|---|---|
| ACAN | Aggrecan | ECM protein | Extracellular matrix | Major NP proteoglycan |
| ADAMTS4/5 | A disintegrin and metalloproteinase with thrombospondin motifs 4 and 5 | Protease | ECM degradation | Aggrecanases |

*Continued on next page*

**Table S1.** Definition of nodes included in the NP mechanotransduction regulatory network (continued).

| Node ID | Full name | Node type | Functional category | Notes |
|---|---|---|---|---|
| AKT1-E | AKT serine/threonine kinase 1 | Kinase | Survival / metabolism | Hyper-loading state variable |
| AKT1-M | AKT serine/threonine kinase 1 | Kinase | Survival / metabolism | Normal-loading state variable |
| AMPK | AMP-activated protein kinase | Kinase | Energy sensing | Catabolic restraint / stress adaptation |
| ANXA1 | Annexin A1 | Signaling protein | Anti-inflammatory | Modulates inflammatory signaling |
| AP-1 | Activator protein 1 (FOS/JUN complex) | Transcription factor | Stress / inflammation | MAPK-responsive TF |
| AQP1 | Aquaporin 1 | Channel | Osmoregulation | Water transport |
| AQP5 | Aquaporin 5 | Channel | Osmoregulation | Water transport |
| BAD | BCL2-associated agonist of cell death | Apoptosis regulator | Cell death | Pro-apoptotic |
| BAX | BCL2-associated X protein | Apoptosis regulator | Cell death | Mitochondrial apoptosis |
| Bcl2 | B-cell lymphoma 2 | Apoptosis regulator | Cell survival | Anti-apoptotic |
| BMP2 | Bone morphogenetic protein 2 | Growth factor | Differentiation / repair | BMP family ligand |
| BNIP3 | BCL2 interacting protein 3 | Stress protein | Mitophagy / apoptosis | Hypoxia/stress-responsive |
| $Ca^{2+}$os | Osmotically regulated intracellular $Ca^{2+}$ signal | Process | Osmosensing | Mechano-osmotic $Ca^{2+}$ |
| $Ca^{2+}$su | Strain-induced intracellular $Ca^{2+}$ signal | Process | Mechanosensing | Strain-driven $Ca^{2+}$ |
| CaMKII | $Ca^{2+}$/calmodulin-dependent protein kinase II | Kinase | $Ca^{2+}$ signaling | Downstream of $Ca^{2+}$ influx |
| CaN | Calcineurin (protein phosphatase 2B/PP2B) | Phosphatase | $Ca^{2+}$ signaling | Activates NFAT |
| CASP3 | Caspase 3 | Protease | Apoptosis | Executioner caspase |
| CASP9 | Caspase 9 | Protease | Apoptosis | Initiator (intrinsic) caspase |
| CAT | Catalase | Enzyme | Antioxidant | $H_2O_2$ detoxification |
| CCL2 | C-C motif chemokine ligand 2 (MCP-1) | Chemokine | Inflammation | Immune recruitment |
| CCN2 | Cellular communication network factor 2 (CTGF) | Matricellular factor | ECM remodeling | Mechanosensitive profibrotic mediator |
| CDC42 | Cell division cycle 42 | Small GTPase | Cytoskeleton | Rho-family GTPase |
| CITED2 | CBP/p300-interacting transactivator with ED-rich tail 2 | Transcriptional regulator | Stress adaptation | Cofactor / transcriptional modulation |
| COL10A1 | Collagen type X alpha 1 chain | ECM protein | Hypertrophy | Hypertrophic marker |
| COL1A1 | Collagen type I alpha 1 chain | ECM protein | Fibrosis | Fibrocartilaginous |
| COL2A1 | Collagen type II alpha 1 chain | ECM protein | NP phenotype | Anabolic marker |
| COX-2 | Cyclooxygenase 2 (PTGS2) | Enzyme | Inflammation | Prostaglandin synthesis |
| CREB | cAMP response element-binding protein | Transcription factor | Survival / metabolism | Activity-dependent TF |
| CXCL1 | C-X-C motif chemokine ligand 1 (GRO-$\alpha$) | Chemokine | Inflammation | Neutrophil chemoattractant |
| CXCL3 | C-X-C motif chemokine ligand 3 (GRO-$\gamma$) | Chemokine | Inflammation | Neutrophil chemoattractant |
| DAG | Diacylglycerol | Second messenger | Lipid signaling | PLC product; activates PKC |

*Continued on next page*

**Table S1.** Definition of nodes included in the NP mechanotransduction regulatory network (continued).

| Node ID | Full name | Node type | Functional category | Notes |
|---|---|---|---|---|
| DRP1 | Dynamin-related protein 1 | GTPase | Mitochondrial dynamics | Mitochondrial fission |
| ELK1 | ETS-like transcription factor 1 | Transcription factor | MAPK signaling | ERK-responsive TF |
| ERK-E | Extracellular signal-regulated kinase 1/2 | Kinase | MAPK signaling | Hyper-loading state variable |
| ERK-M | Extracellular signal-regulated kinase 1/2 | Kinase | MAPK signaling | Normal-loading state variable |
| FAK-E | Focal adhesion kinase (PTK2) | Kinase | Mechanotransduction | Hyper-loading state variable |
| FAK-M | Focal adhesion kinase (PTK2) | Kinase | Mechanotransduction | Normal-loading state variable |
| FGF18 | Fibroblast growth factor 18 | Growth factor | Growth / repair | Pro-regenerative factor |
| FGF2 | Fibroblast growth factor 2 (basic FGF) | Growth factor | Growth / repair | Mitogenic factor |
| FOXO | Forkhead box O transcription factors (FOXO family) | Transcription factor | Stress resistance | Antioxidant/autophagy programs |
| GADD45 | Growth arrest and DNA damage-inducible 45 | Stress protein | DNA damage response | Stress-induced regulator |
| GDF5 | Growth differentiation factor 5 | Growth factor | Cartilage homeostasis | BMP family member |
| GPX1 | Glutathione peroxidase 1 | Enzyme | Antioxidant | Peroxide detoxification |
| GSK3B | Glycogen synthase kinase 3 beta | Kinase | Wnt/metabolism | Regulates $\beta$-catenin, FOXO, etc. |
| HIF-1$\alpha$ | Hypoxia-inducible factor 1 alpha | Transcription factor | Hypoxia | Adaptive hypoxia signaling |
| HIF-2$\alpha$ | Hypoxia-inducible factor 2 alpha (EPAS1) | Transcription factor | Hypoxia / degeneration | Often linked to catabolic bias |
| HL | Hyper mechanical loading | External input | Mechanical stimulus | Excessive loading input |
| HO-1 | Heme oxygenase 1 (HMOX1) | Enzyme | Antioxidant | Nrf2 target; cytoprotection |
| HSF1 | Heat shock factor 1 | Transcription factor | Proteostasis | Heat-shock response TF |
| HSP27 | Heat shock protein 27 (HSPB1) | Chaperone | Proteostasis | Small HSP |
| HSP70 | Heat shock protein 70 (HSPA family) | Chaperone | Proteostasis | Stress chaperone |
| IGF1 | Insulin-like growth factor 1 | Growth factor | Growth / survival | Pro-anabolic signaling |
| IKK | I$\kappa$B kinase complex | Kinase complex | Inflammation | Activates NF-$\kappa$B |
| IL1$\beta$ | Interleukin 1 beta | Cytokine | Inflammation | Pro-inflammatory cytokine |
| IL6 | Interleukin 6 | Cytokine | Inflammation | Pro-inflammatory cytokine |
| IL8 | Interleukin 8 (CXCL8) | Chemokine | Inflammation | Neutrophil chemoattractant |
| iNOS | Inducible nitric oxide synthase (NOS2) | Enzyme | Inflammation / nitrosative stress | Produces nitric oxide |
| IP3 | Inositol 1,4,5-trisphosphate | Second messenger | $Ca^{2+}$ signaling | ER $Ca^{2+}$ release |
| I$\kappa$B$\alpha$ | Inhibitor of NF-$\kappa$B alpha (NFKBIA) | Inhibitory protein | Inflammation | Sequesters NF-$\kappa$B |
| JNK | c-Jun N-terminal kinase (JNK family) | Kinase | MAPK signaling | Stress-activated MAPK |
| LATS1/2 | Large tumor suppressor kinases 1 and 2 | Kinase | Hippo signaling | YAP/TAZ inhibition |
| LIMK1 | LIM domain kinase 1 | Kinase | Cytoskeleton | Actin remodeling |

*Continued on next page*

**Table S1.** Definition of nodes included in the NP mechanotransduction regulatory network (continued).

| Node ID | Full name | Node type | Functional category | Notes |
|---|---|---|---|---|
| MAP3K | Mitogen-activated protein kinase kinase kinase (generic) | Kinase | MAPK signaling | Upstream MAPK activator |
| MEK-E | MAPK/ERK kinase 1/2 (MEK1/2) | Kinase | MAPK signaling | Hyper-loading state variable |
| MEK-M | MAPK/ERK kinase 1/2 (MEK1/2) | Kinase | MAPK signaling | Normal-loading state variable |
| MKK3/6 | MAPK kinase 3 and 6 | Kinase | MAPK signaling | Activates p38 |
| MKK4/7 | MAPK kinase 4 and 7 | Kinase | MAPK signaling | Activates JNK |
| MMP1 | Matrix metalloproteinase 1 | Protease | ECM degradation | Collagenase-1 |
| MMP13 | Matrix metalloproteinase 13 | Protease | ECM degradation | Collagenase-3 |
| MMP3 | Matrix metalloproteinase 3 | Protease | ECM degradation | Stromelysin-1 |
| MOMP | Mitochondrial outer membrane permeabilization | Process | Apoptosis | Intrinsic apoptosis commitment |
| MRTF-A | Myocardin-related transcription factor A (MKL1) | Transcriptional cofactor | Cytoskeleton-linked TF | SRF/MRTF axis |
| MST1/2 | Mammalian sterile 20-like kinases 1 and 2 | Kinase | Hippo signaling | Upstream of LATS1/2 |
| mTORC1 | Mechanistic target of rapamycin complex 1 | Kinase complex | Metabolism / growth | Protein synthesis, autophagy inhibition |
| mTORC2 | Mechanistic target of rapamycin complex 2 | Kinase complex | Metabolism / survival | AKT regulation, cytoskeleton |
| NF-$\kappa$B | Nuclear factor kappa B | Transcription factor | Inflammation | Typically RELA/p50 |
| NFAT | Nuclear factor of activated T cells (NFATc family) | Transcription factor | $Ca^{2+}$ signaling | Calcineurin-dependent (NFATc1–c4) |
| NL | Normal mechanical loading | External input | Mechanical stimulus | Physiological loading input |
| NRF2 | Nuclear factor erythroid 2–related factor 2 (NFE2L2) | Transcription factor | Antioxidant | Master redox regulator |
| P38 | p38 mitogen-activated protein kinase (p38 MAPK) | Kinase | MAPK signaling | Stress-activated MAPK |
| PAK1 | p21-activated kinase 1 | Kinase | Cytoskeleton | RAC/CDC42 effector |
| PDK1-E | 3-phosphoinositide-dependent protein kinase 1 | Kinase | PI3K/AKT signaling | Hyper-loading state variable |
| PDK1-M | 3-phosphoinositide-dependent protein kinase 1 | Kinase | PI3K/AKT signaling | Normal-loading state variable |
| PGE2 | Prostaglandin $E_2$ | Lipid mediator | Inflammation | COX-2 product |
| PHD2 | Prolyl hydroxylase domain-containing protein 2 (EGLN1) | Enzyme | Hypoxia sensing | Regulates HIF stability |
| PI3K-E | Phosphoinositide 3-kinase (class I PI3K) | Kinase | PI3K/AKT signaling | Hyper-loading state variable |
| PI3K-M | Phosphoinositide 3-kinase (class I PI3K) | Kinase | PI3K/AKT signaling | Normal-loading state variable |
| PIEZO1 | PIEZO-type mechanosensitive ion channel component 1 | Ion channel | Mechanosensing | Strain-sensitive $Ca^{2+}$ entry |
| PIP3-E | Phosphatidylinositol (3,4,5)-trisphosphate | Lipid messenger | PI3K signaling | Hyper-loading state variable |
| PIP3-M | Phosphatidylinositol (3,4,5)-trisphosphate | Lipid messenger | PI3K signaling | Normal-loading state variable |
| PKC-M | Protein kinase C (PKC family) | Kinase | DAG signaling | Normal-loading state variable |

*Continued on next page*

**Table S1.** Definition of nodes included in the NP mechanotransduction regulatory network (continued).

| Node ID | Full name | Node type | Functional category | Notes |
|---|---|---|---|---|
| PKC-E | Protein kinase C (PKC family) | Kinase | DAG signaling | Hyper-loading state variable |
| PKN1 | Protein kinase N1 (PRK1) | Kinase | Cytoskeleton | RhoA effector kinase |
| PLA2 | Phospholipase $A_2$ ($PLA_2$ family) | Enzyme | Lipid signaling | Releases arachidonic acid |
| PLC$\gamma$-M | Phospholipase C gamma (PLC$\gamma$ family) | Enzyme | $Ca^{2+}$/DAG signaling | Normal-loading state variable |
| PLC$\gamma$-E | Phospholipase C gamma (PLC$\gamma$ family) | Enzyme | $Ca^{2+}$/DAG signaling | Hyper-loading state variable |
| PLD2 | Phospholipase D2 | Enzyme | Lipid signaling | Produces phosphatidic acid |
| PPAR$\gamma$ | Peroxisome proliferator-activated receptor gamma | Nuclear receptor | Metabolism / inflammation | Anti-inflammatory metabolic regulator |
| PTEN | Phosphatase and tensin homolog | Phosphatase | PI3K/AKT regulation | Antagonizes PIP3/AKT |
| RAC1-M | Ras-related C3 botulinum toxin substrate 1 | Small GTPase | Cytoskeleton | Normal-loading state variable |
| RAC1-E | Ras-related C3 botulinum toxin substrate 1 | Small GTPase | Cytoskeleton | Hyper-loading state variable |
| RAF-E | RAF serine/threonine kinase (RAF family) | Kinase | MAPK signaling | Hyper-loading state variable |
| RAF-M | RAF serine/threonine kinase (RAF family) | Kinase | MAPK signaling | Normal-loading state variable |
| RAS-E | RAS small GTPase (RAS family) | Small GTPase | MAPK/PI3K entry | Hyper-loading state variable |
| RAS-M | RAS small GTPase (RAS family) | Small GTPase | MAPK/PI3K entry | Normal-loading state variable |
| Rheb | Ras homolog enriched in brain | Small GTPase | mTOR signaling | Activates mTORC1 |
| RhoA-E | Ras homolog family member A | Small GTPase | Cytoskeleton | Hyper-loading state variable |
| RhoA-M | Ras homolog family member A | Small GTPase | Cytoskeleton | Normal-loading state variable |
| ROCK-E | Rho-associated coiled-coil-containing protein kinase (ROCK1/2) | Kinase | Cytoskeleton | Hyper-loading state variable |
| ROCK-M | Rho-associated coiled-coil-containing protein kinase (ROCK1/2) | Kinase | Cytoskeleton | Normal-loading state variable |
| ROS | Reactive oxygen species | Process | Oxidative stress | Redox signaling/oxidative load |
| RSK | Ribosomal S6 kinase (p90RSK) | Kinase | MAPK signaling | ERK downstream effector |
| RUNX2 | Runt-related transcription factor 2 | Transcription factor | Hypertrophy / ossification | Fibrocartilaginous/hypertrophic TF |
| SIRT1 | Sirtuin 1 | Deacetylase | Metabolism / stress | $NAD^{+}$-dependent deacetylase |
| SMIT1 | Sodium/myo-inositol cotransporter 1 (SLC5A3) | Transporter | Osmoregulation | Osmolyte transporter |
| SOD1 | Superoxide dismutase 1 | Enzyme | Antioxidant | Cytosolic superoxide detoxification |
| SOD2 | Superoxide dismutase 2 | Enzyme | Antioxidant | Mitochondrial superoxide detoxification |
| SOX5 | SRY-box transcription factor 5 | Transcription factor | NP/chondrogenic phenotype | Chondrogenic regulator |

*Continued on next page*

**Table S1.** Definition of nodes included in the NP mechanotransduction regulatory network (continued).

| Node ID | Full name | Node type | Functional category | Notes |
|---|---|---|---|---|
| SOX6 | SRY-box transcription factor 6 | Transcription factor | NP/chondrogenic phenotype | Chondrogenic regulator |
| SOX9 | SRY-box transcription factor 9 | Transcription factor | NP/chondrogenic phenotype | Master anabolic regulator |
| TAK1 | TGF-$\beta$-activated kinase 1 (MAP3K7) | Kinase | Inflammation / MAPK | Upstream of NF-$\kappa$B/JNK/p38 |
| TAUT | Taurine transporter (SLC6A6) | Transporter | Osmoregulation | Osmolyte transport |
| TGF$\beta$ | Transforming growth factor beta (TGF-$\beta$ family) | Growth factor | Anabolism / fibrosis | Includes latent activation contexts |
| TIMP3 | Tissue inhibitor of metalloproteinases 3 | Inhibitor | ECM protection | Inhibits MMP/ADAMTS |
| TNF | Tumor necrosis factor (TNF-$\alpha$) | Cytokine | Inflammation | Pro-inflammatory cytokine |
| TonEBP | Tonicity-responsive enhancer-binding protein (NFAT5) | Transcription factor | Osmoregulation | Osmoadaptive TF |
| TRPV4 | Transient receptor potential vanilloid 4 | Ion channel | Osmomechanosensing | Mechanosensitive $Ca^{2+}$ channel |
| ULK1 | Unc-51-like autophagy activating kinase 1 | Kinase | Autophagy | Initiates autophagy |
| VEGF | Vascular endothelial growth factor (VEGF-A) | Growth factor | Angiogenesis | Degeneration-associated angiogenic cue |
| VHL | Von Hippel–Lindau tumor suppressor | E3 ligase component | Hypoxia sensing | Targets HIF for degradation |
| Wnt3a | Wnt family member 3A | Growth factor (ligand) | Wnt signaling | Canonical Wnt ligand |
| Wnt5a | Wnt family member 5A | Growth factor (ligand) | Wnt signaling | Non-canonical Wnt ligand |
| YAP/TAZ | Yes-associated protein / transcriptional co-activator with PDZ-binding motif (TAZ/WWTR1) | Transcriptional co-activator | Mechanotransduction | Hippo/cytoskeleton-linked |
| $\alpha_5\beta_1$-FN | $\alpha_5\beta_1$ integrin (intact fibronectin-binding configuration) | Integrin receptor | Mechanosensing / adhesion | Homeostatic ECM context |
| $\alpha_5\beta_1$-Fs | $\alpha_5\beta_1$ integrin (fragmented fibronectin-binding configuration) | Integrin receptor | Mechanosensing / adhesion | Degenerative ECM context |
| $\alpha_V\beta_3$ | $\alpha_V\beta_3$ integrin | Integrin receptor | Mechanosensing / adhesion | Strain/inflammatory bias |
| $\alpha_V\beta_6$ | $\alpha_V\beta_6$ integrin | Integrin receptor | TGF-$\beta$ activation | Activates latent TGF-$\beta$ |
| $\beta$-catenin | Beta-catenin (CTNNB1) | Transcriptional co-activator | Wnt signaling | Canonical Wnt effector |
| $NAD^+$ | Nicotinamide adenine dinucleotide (oxidized form) | Metabolite | Redox / metabolism | SIRT1 substrate/cofactor |
| LKB1 | Liver kinase B1 (STK11) | Kinase | Energy sensing | Upstream AMPK kinase |
| SDC4-M | Syndecan-4 | Proteoglycan receptor | Mechanosensing | Normal-loading state variable |
| SDC4-E | Syndecan-4 | Proteoglycan receptor | Mechanosensing | Hyper-loading state variable |
| Hypo | Hypo-loading | Mechanical input | Mechanotransduction reduction | Associated with impaired cellular signaling and matrix homeostasis |
| MitD | Mitochondrial dysfunction | Cellular condition | Energy metabolism disruption | Associated with reduced ATP production and increased oxidative stress |

#### Knowledge-driven RNM construction and curation workflow

The overall workflow used to construct, analyse, and falsify the knowledge-driven RNM is summarized in Fig. S1. The pipeline integrates systematic literature curation, network topology construction, dynamical modelling, and iterative falsification and refinement steps to derive a predictive regulatory framework for nucleus pulposus (NP) cell responses to mechanical loading.

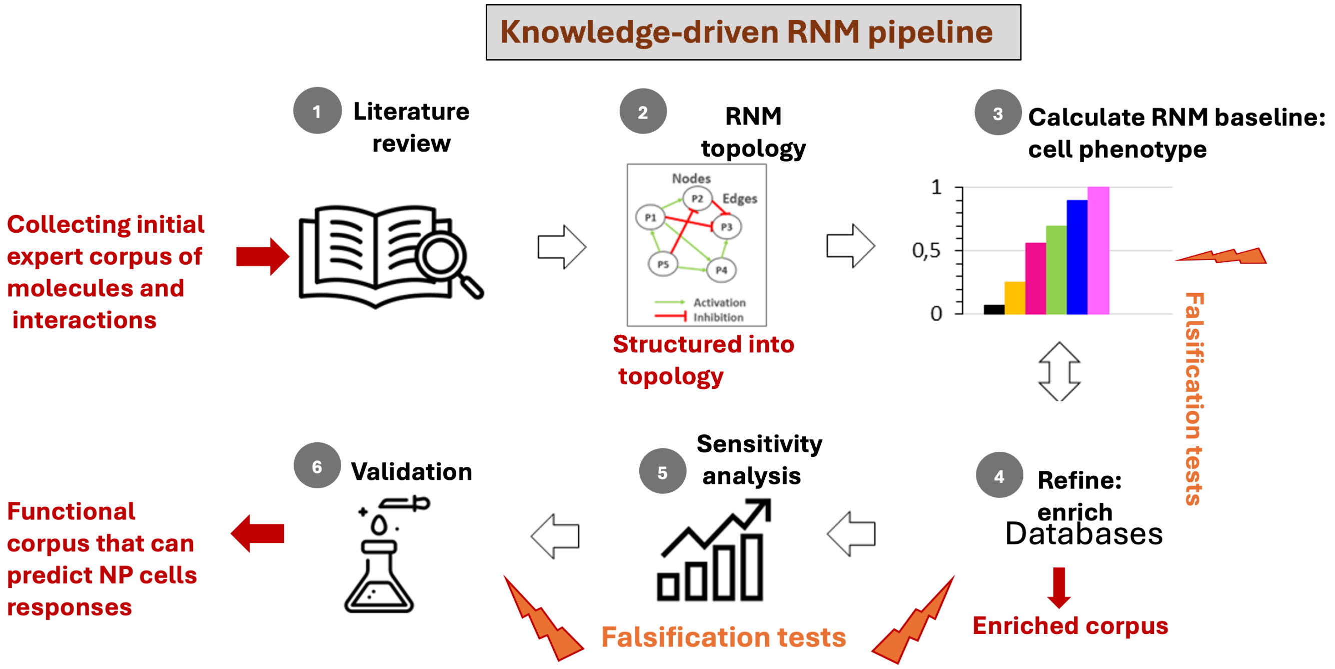


**Figure S1.** Knowledge-driven regulatory network model (RNM) construction and falsification pipeline. Mechanotransduction knowledge was first collected through structured literature review and expert curation of experimentally supported molecular interactions. These interactions were organized into a directed regulatory network topology describing activation and inhibition relationships among nodes. The resulting RNM was formulated as a semi-quantitative dynamical model to compute steady-state cellular phenotypes under different mechanical loading regimes. Model predictions were subjected to falsification tests against independent experimental load–response studies and further analysed using sensitivity analysis. Iterative refinement using pathway databases and additional literature evidence allowed enrichment of the interaction corpus, yielding a falsified functional network capable of predicting NP cell responses to mechanical loading.

**Literature search strategy** Mechanotransductive interactions were curated using structured literature searches across PubMed, Scopus, Web of Science, and Google Scholar, following established guidelines for reproducible evidence synthesis[1,2]. Searches employed Boolean operators and controlled vocabulary and were iteratively refined to ensure coverage of NP-specific and closely related mechanobiological studies[3,4]. The following search strings were used:

- (**Search 1**) (“nucleus pulposus” OR “intervertebral disc”) AND (“mechanotransduction” OR “mechanical loading”)
- (**Search 2**) (“nucleus pulposus” OR “chondrocyte”) AND (“mechanotransduction” OR “mechanical loading”)
- (**Search 3**) (“nucleus pulposus” OR “intervertebral disc”) AND (“mechanobiology” OR “mechanical loading”)
- (**Search 4**) (“nucleus pulposus” OR “intervertebral disc”) AND (“mechanobiology” OR “mechanical loading”) AND (“Node name”)

Inclusion criteria required direct experimental evidence of mechanical modulation of molecular activity. When NP-specific evidence was unavailable, mechanosensitive interactions from articular cartilage, mesenchymal stem cells, or fibrocartilaginous tissues were included if regulatory directionality was conserved across systems[5,6].

**Network encoding and node inventory:** All curated interactions were encoded as activating (+) or inhibitory (−) edges in a directed signed graph[7,8]. The network includes mechanical stimuli; primary mechanosensors ($\alpha_5\beta_1$, $\alpha_V\beta_3$, $\alpha_V\beta_6$, PIEZO1, TRPV4); cytoskeletal regulators (RhoA/ROCK); kinase cascades (FAK, MAPK/ERK, JNK, p38, PI3K/AKT/mTOR); redox and inflammatory regulators (NRF2/ROS, NF-$\kappa$B, AP-1); growth factors (TGF-$\beta$, BMPs, IGF-1, GDF5); transcription factors (SOX9, RUNX2, FOXO, CREB, YAP/TAZ, TonEBP, HIFs); extracellular matrix effectors (*COL2A1*, *ACAN*, *MMPs*, *ADAMTS*); apoptosis and stress-response mediators (BCL2, BAX, CASP3, CASP9, BNIP3, NIX, GADD45); and oxidative-stress/proteostasis regulators (HO-1, GPX1, SOD1/2, CAT, HSP70, HSP27)[9].

#### Complementary node-resolved and phenotype-level analyses of the rescue screen

The perturbation screen was further characterized at the individual-node level and by evaluating movement of a broader anabolic/catabolic phenotype toward the Normal-loading state. Both analyses used the replicate-specific Hyper-to-perturbation trajectories described in the main Methods.

##### *Paired node-level perturbation responses*

Because each perturbation simulation was initialized from its corresponding Hyper steady-state replicate, intervention-induced changes in individual nodes were evaluated on a paired basis. For node $i$, perturbation $p$, and replicate $r$, the perturbation response was defined as

$$\Delta_{r,i}^{p} = x_{r,i}^{p} - x_{r,i}^{\text{Hyper}}, \tag{1}$$

where $x_{r,i}^{\text{Hyper}}$ and $x_{r,i}^{p}$ denote the steady-state activities of the same node before and after perturbation, respectively. The mean response across the $R = 100$ paired replicates was

$$\overline{\Delta}_{i}^{p} = \frac{1}{R}\sum_{r=1}^{R} \Delta_{r,i}^{p}, \tag{2}$$

with the corresponding standard deviation

$$s_{\Delta,i}^{p} = \left[\frac{1}{R-1}\sum_{r=1}^{R}\left(\Delta_{r,i}^{p} - \overline{\Delta}_{i}^{p}\right)^{2}\right]^{1/2}. \tag{3}$$

Positive values of $\overline{\Delta}_{i}^{p}$ indicate increased steady-state activity relative to the Hyper-loading state, whereas negative values indicate decreased activity. For each functional module $\mathscr{C}$, the overall magnitude of the node-level response to perturbation $p$ was summarized as

$$M_{p,\mathscr{C}} = \frac{1}{|\mathscr{C}|}\sum_{i\in\mathscr{C}} \left|\overline{\Delta}_{i}^{p}\right|, \tag{4}$$

where $|\mathscr{C}|$ denotes the number of nodes in the module. Interventions were ranked according to $M_{p,\mathscr{C}}$ to identify the 10 perturbations producing the largest overall node-level response within each module. A leading dual-node intervention among these top-ranked perturbations and the highest-ranked single-node intervention were further examined at the individual-node level. Because $M_{p,\mathscr{C}}$ quantifies response magnitude without reference to the Normal-loading state, it is distinct from the Hyper-to-Normal rescue percentage used for functional-module restoration.

##### *Phenotype-level distance to the Normal-loading state*

A complementary analysis evaluated whether each perturbation shifted a predefined anabolic/catabolic phenotype toward the Normal-loading state at the replicate level. The anabolic readout panel comprised ACAN, COL2A1, TIMP3, PPAR$\gamma$, GDF5, IGF1, SOX9, and Bcl2, whereas the catabolic panel comprised MMP1, MMP3, MMP13, ADAMTS4/5, IL1$\beta$, IL6, IL8, TNF, iNOS, and COX-2. Let $\mathscr{S}$ denote the union of these readouts.

For Hyper replicate $r$, the Euclidean distance to the ensemble-mean Normal-loading reference profile was

$$d_{r}^{\text{Hyper}} = \left\| \mathbf{x}_{r,\mathscr{S}}^{\text{Hyper}} - \bar{\mathbf{x}}_{\mathscr{S}}^{\text{Normal}} \right\|_{2}, \tag{5}$$

and the corresponding distance following perturbation $p$ was

$$d_{r}^{p} = \left\| \mathbf{x}_{r,\mathscr{S}}^{p} - \bar{\mathbf{x}}_{\mathscr{S}}^{\text{Normal}} \right\|_{2}. \tag{6}$$

The perturbation-induced change in mean distance to the Normal-loading phenotype was calculated as

$$\Delta D_{p} = \frac{1}{R}\sum_{r=1}^{R} d_{r}^{p} - \frac{1}{R}\sum_{r=1}^{R} d_{r}^{\text{Hyper}}. \tag{7}$$

Accordingly, $\Delta D_p < 0$ indicates that perturbation $p$ reduced the average distance of the anabolic/catabolic phenotype from the Normal-loading reference, whereas $\Delta D_p > 0$ indicates increased separation from Normal. This analysis provides a replicate-level measure of broader phenotypic movement toward the Normal state and complements both the node-resolved response analysis and the functional-module rescue metric described in the main Methods.

### 2 Results

#### Model falsification test

The complete node-level results of the normal-loading (NL) versus hyperphysiological-loading (HL) comparison are provided in Table S2. For each of the 39 retained literature-derived benchmarks, the table reports the literature-derived response direction (HL-like $\rightarrow$ NL-like), mean final RNM activation under NL and HL (100 replicate simulations per condition), the loading effect ($\Delta_i = \bar{x}_i^{\mathrm{NL}} - \bar{x}_i^{\mathrm{HL}}$), its 95% percentile bootstrap confidence interval (CI), and the directional outcome. An upward arrow indicates higher expression or activity under NL-like conditions, and a downward arrow indicates higher expression or activity under HL-like conditions. The benchmarks comprise 6 upward-response and 33 downward-response markers.

Using the CI-based criteria defined in the Methods, 37 of 39 predictions (94.9%) were concordant with their literature-derived response directions, including 6/6 + literature-derived response and 31/33 − literature-derived response benchmarks. HSF1 and CCN2 were contradictory, and none was indeterminate. The bootstrap CIs quantify uncertainty in the estimated model effects arising from the replicate simulation ensembles.

**Table S2.** RNM loading effects and 95% bootstrap confidence intervals for the 39 retained NP mechanobiology benchmarks. Numerical values are rounded to five decimal places.

| Node | Lit. dir. | Mean NL | Mean HL | $\Delta_i$ | 95% bootstrap CI | Outcome |
|---|---|---|---|---|---|---|
| *+ literature-derived response (6 nodes)* | | | | | | |
| ACAN | + | 0.99856 | 0.18315 | +0.81541 | [+0.74881, +0.88201] | Concordant |
| Bcl2 | + | 0.99249 | 0.00002 | +0.99247 | [+0.99217, +0.99277] | Concordant |
| COL2A1 | + | 0.99849 | 0.21336 | +0.78513 | [+0.70755, +0.86272] | Concordant |
| SOD2 | + | 0.91006 | 0.20751 | +0.70254 | [+0.62213, +0.78069] | Concordant |
| SOX9 | + | 0.99941 | 0.00006 | +0.99934 | [+0.99929, +0.99939] | Concordant |
| TIMP3 | + | 0.99825 | 0.00000 | +0.99825 | [+0.99796, +0.99854] | Concordant |
| *− literature-derived response (33 nodes)* | | | | | | |
| ADAMTS4/5 | − | 0.00000 | 0.99845 | -0.99845 | [-0.99901, -0.99788] | Concordant |
| BAX | − | 0.00000 | 0.78000 | -0.78000 | [-0.86000, -0.70000] | Concordant |
| CASP3 | − | 0.00000 | 0.78000 | -0.78000 | [-0.86000, -0.70000] | Concordant |
| CASP9 | − | 0.00000 | 0.78000 | -0.78000 | [-0.86000, -0.70000] | Concordant |
| CCL2 | − | 0.00000 | 0.99980 | -0.99980 | [-0.99987, -0.99973] | Concordant |
| CCN2 | − | 0.98427 | 0.64413 | +0.34014 | [+0.27411, +0.41441] | **Contradictory** |
| COL10A1 | − | 0.00000 | 0.98354 | -0.98354 | [-0.98947, -0.97760] | Concordant |
| COL1A1 | − | 0.00119 | 0.99946 | -0.99827 | [-0.99849, -0.99806] | Concordant |
| COX-2 | − | 0.00000 | 0.99990 | -0.99990 | [-0.99993, -0.99986] | Concordant |
| CXCL1 | − | 0.00000 | 0.99980 | -0.99980 | [-0.99987, -0.99973] | Concordant |
| DRP1 | − | 0.00000 | 0.97785 | -0.97785 | [-0.98590, -0.96980] | Concordant |
| GSK3B | − | 0.00000 | 0.78000 | -0.78000 | [-0.86000, -0.70000] | Concordant |
| HSF1 | − | 0.51464 | 0.02747 | +0.48718 | [+0.39947, +0.57455] | **Contradictory** |
| IL1$\beta$ | − | 0.00317 | 0.99946 | -0.99628 | [-0.99685, -0.99570] | Concordant |
| IL6 | − | 0.00317 | 0.99946 | -0.99628 | [-0.99686, -0.99571] | Concordant |
| IL8 | − | 0.00000 | 0.99980 | -0.99980 | [-0.99987, -0.99973] | Concordant |
| JNK | − | 0.00004 | 0.99699 | -0.99695 | [-0.99805, -0.99586] | Concordant |
| MMP13 | − | 0.00001 | 0.99947 | -0.99946 | [-0.99961, -0.99930] | Concordant |
| MMP3 | − | 0.00001 | 0.99947 | -0.99946 | [-0.99950, -0.99942] | Concordant |
| MRTF-A | − | 0.47045 | 0.98493 | -0.51448 | [-0.60091, -0.42872] | Concordant |
| NF-$\kappa$B | − | 0.00000 | 0.98375 | -0.98375 | [-0.98966, -0.97784] | Concordant |
| P38 | − | 0.00004 | 0.99699 | -0.99695 | [-0.99805, -0.99586] | Concordant |
| PGE2 | − | 0.83346 | 0.99993 | -0.16647 | [-0.16647, -0.16647] | Concordant |
| PIEZO1 | − | 0.00144 | 0.98263 | -0.98119 | [-0.98119, -0.98119] | Concordant |
| PKN1 | − | 0.00001 | 1.00000 | -0.99999 | [-0.99999, -0.99999] | Concordant |
| ROS | − | 0.00001 | 0.78000 | -0.77999 | [-0.85999, -0.69999] | Concordant |
| RUNX2 | − | 0.00001 | 0.98634 | -0.98633 | [-0.98637, -0.98629] | Concordant |
| TGF$\beta$ | − | 0.84641 | 0.98594 | -0.13953 | [-0.13953, -0.13953] | Concordant |
| TNF | − | 0.00000 | 0.99990 | -0.99990 | [-0.99993, -0.99986] | Concordant |
| VEGF | − | 0.77248 | 0.98571 | -0.21323 | [-0.22602, -0.20038] | Concordant |
| Wnt3a | − | 0.00001 | 0.98876 | -0.98875 | [-0.98979, -0.98770] | Concordant |
| YAP/TAZ | − | 0.82450 | 0.92998 | -0.10548 | [-0.10548, -0.10548] | Concordant |
| $\beta$-catenin | − | 0.04891 | 0.25635 | -0.20744 | [-0.28209, -0.13280] | Concordant |

#### Hyper-to-Normal rescue screening via node clamping

**Functional-module rescue rankings.** The highest-scoring intervention for different functional modules is listed in Table S3. Rescue percentages refer to reductions in Euclidean distance to the NL ensemble-mean module profile relative to the corresponding unperturbed HL-to-NL distance.

**Table S3.** Highest-scoring intervention for different functional modules.

| Functional module | Top-ranked intervention | Rescue (%) |
|---|---|---|
| ECM anabolism and phenotype markers | SOX9-ROS | 85.9 |
| Oxidative-stress defense and proteostasis | FOXO-ROS | 68.9 |
| Growth factors | SOX9-NF-$\kappa$B | 66.0 |
| Cytokines, chemokines, proteases, and others | SOX9-NF-$\kappa$B | 57.0 |
| Cell survival, apoptosis, mitophagy, and DNA damage | NRF2-PIEZO1 | 37.5 |
| Transcription factors | SOX9-ROS | 30.7 |
| Metabolic and related | PPAR$\gamma$-ROS | 23.3 |
| Ion channels and related | PPAR$\gamma$-PIEZO1 | 10.4 |

**Node-resolved intervention responses.** Figs. S2 and S3 show node-level responses for eight selected modules. Within each module, the two interventions with the highest module-level rescue scores were selected from the 48 screened strategies. Points show mean steady-state activation differences between each intervention and its paired HL replicate; error bars show the standard deviation of these differences across 100 simulations. Complete node-level values for all 48 interventions and all 11 modules will be provided as Supplementary Data S1.

**Phenotype-level readouts.** Table S4 reports mean activation of the predefined eight-marker anabolic and ten-marker catabolic panels for interventions ranked among the top ten in either the ECM or cytokine/chemokine/protease module. These selections comprise 12 distinct interventions.

**Table S4.** Mean anabolic- and catabolic-panel activation under NL, HL, and interventions ranked among the top ten in the ECM or cytokine/chemokine/protease module. The union of the two Top-10 lists comprises 12 distinct interventions.

| Regime / intervention | ECM rank | Cytokine rank | Anabolic mean | Catabolic mean |
|---|---|---|---|---|
| NL baseline | – | – | 0.985 | 0.001 |
| HL baseline | – | – | 0.059 | 0.973 |
| SOX9–ROS | 1 | 6 | 0.589 | 0.653 |
| SOX9–NF-$\kappa$B | 2 | 1 | 0.718 | 0.371 |
| SOX9–P38 | 3 | 8 | 0.561 | 0.654 |
| SOX9–mTORC1 | 4 | 10 | 0.562 | 0.655 |
| SOX9–PIEZO1 | 5 | 5 | 0.557 | 0.645 |
| SOX9–RhoA-E | 6 | 3 | 0.556 | 0.628 |
| SOX9 | 7 | – | 0.556 | 0.656 |
| PPAR$\gamma$–ROS | 8 | 9 | 0.463 | 0.655 |
| PPAR$\gamma$–P38 | 9 | – | 0.434 | 0.658 |
| PPAR$\gamma$–PIEZO1 | 10 | 7 | 0.431 | 0.649 |
| PPAR$\gamma$–NF-$\kappa$B | – | 2 | 0.458 | 0.445 |
| PPAR$\gamma$–RhoA-E | – | 4 | 0.430 | 0.634 |

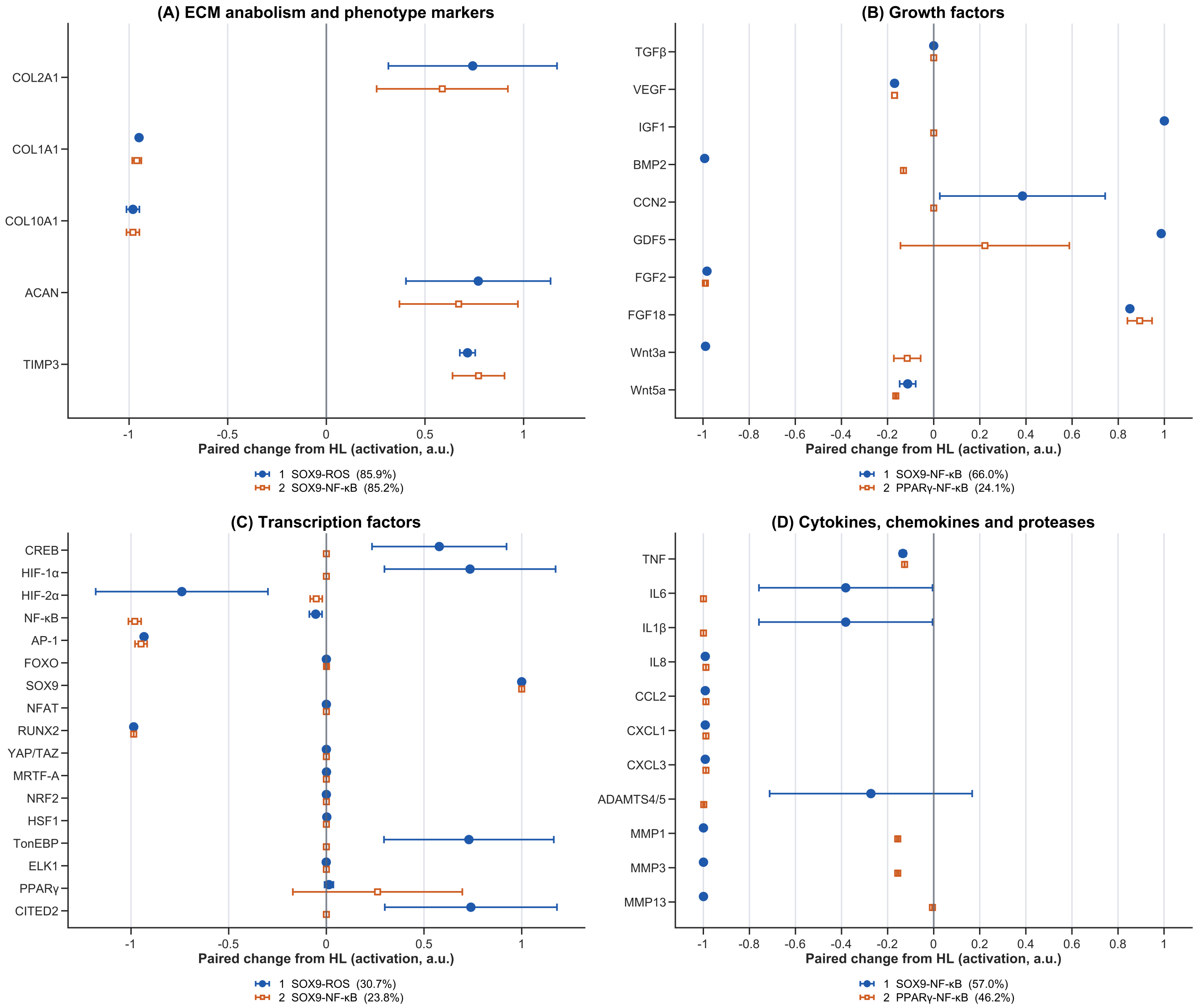


**Figure S2.** Node-resolved responses for the two highest-ranked interventions in four functional modules: (A) ECM anabolism and phenotype markers, (B) growth factors, (C) transcription factors, and (D) cytokines, chemokines, proteases, and others. Blue circles and orange squares identify the first- and second-ranked interventions within each module, respectively. Points indicate mean paired intervention-minus-HL activation changes; horizontal error bars show SD across 100 matched simulations.

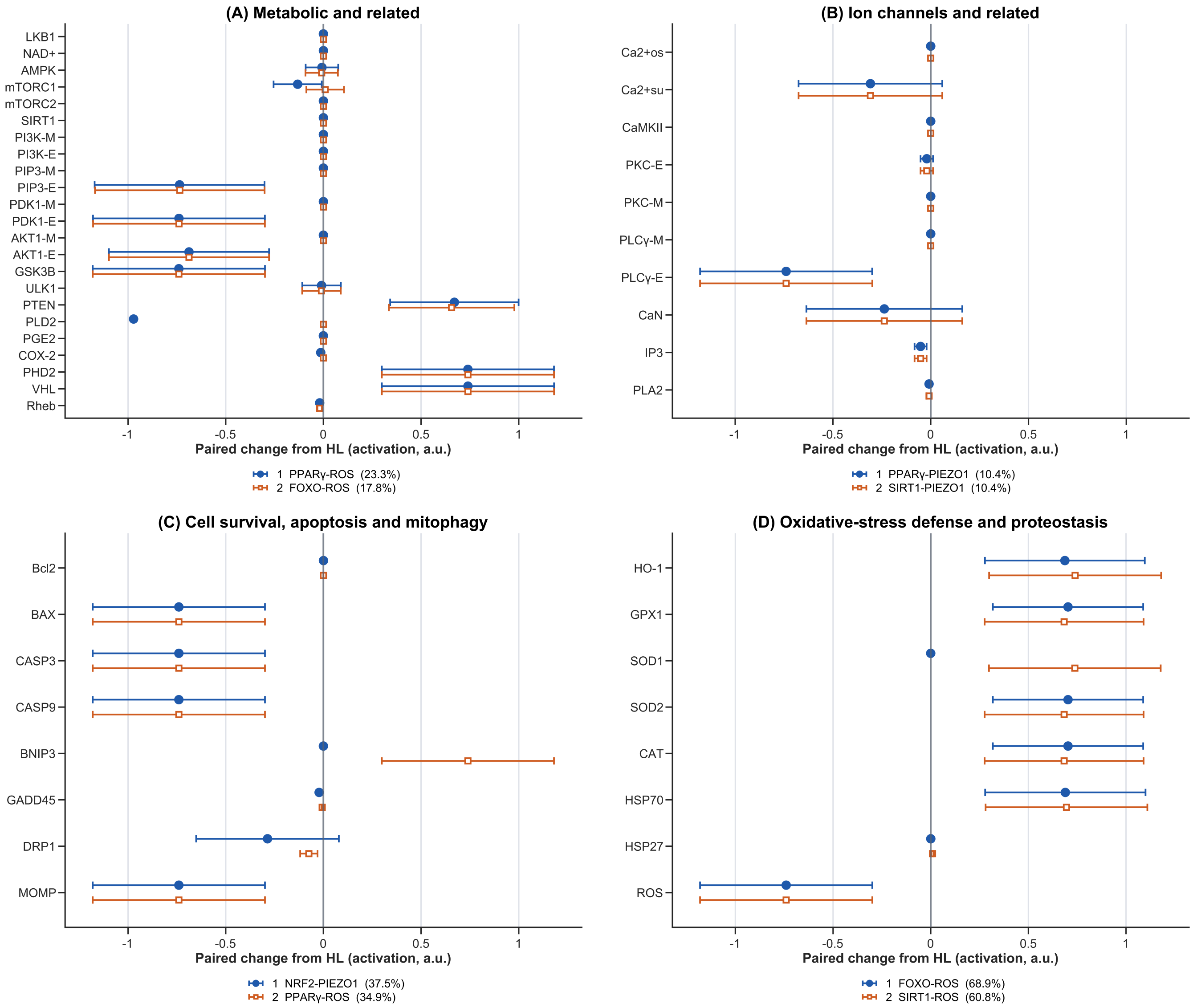


**Figure S3.** Node-resolved responses for the two highest-ranked interventions in four functional modules: (A) metabolic and related regulators, (B) ion channels and related processes, (C) cell survival, apoptosis, mitophagy, and DNA damage, and (D) oxidative-stress defense and proteostasis. Blue circles and orange squares identify the first- and second-ranked interventions within each module, respectively. Points indicate mean paired intervention-minus-HL activation changes; horizontal error bars show SD across 100 matched simulations.
